# Human pancreatic organoids derived from pluripotent stem cells recapitulate pancreatic organogenesis

**DOI:** 10.1101/2025.10.31.685661

**Authors:** Jonathan A. Brassard, Patrizia Tornabene, Daniel O. Kechele, Lily Deng, Julie B. Sneddon, Mansa Krishnamurthy, James M. Wells

## Abstract

Pancreas organogenesis relies on sequential interactions between the pancreatic epithelium and surrounding mesodermal cell types that initiate epithelial budding and branching to form a complex ductal network with terminal acini. Despite recent advances with pluripotent stem cell-based approaches, there are no models that robustly recapitulate pancreas morphogenesis or the spatial organization of ductal, exocrine, endocrine and mesenchymal cells seen in the native organ. Here, we introduce a new pluripotent stem cell-based pancreatic organoid that captures the complexity seen during pancreatic development, with budding and stratification of multipotent progenitors followed by formation of a ductal network that give rise to peripheral acini. We identify a critical role for mesenchyme-derived factors to robustly promote pancreatic organoid formation and morphogenesis. Human pancreatic organoids are correctly patterned, with functional exocrine acini secreting digestive enzymes into a ductal network. Comparative analysis confirms that the pancreatic organoids are similar to early second trimester human pancreas, with potential to further mature upon transplantation in mice. Finally, we show that endocrinogenesis can be reproduced in organoids, generating functional islet-like clusters interspersed within the ductal network. Together, this represents an exciting new platform to study human pancreas development and a broad array of pancreatic diseases.

## Main

The pancreas is a complex organ comprised of endocrine islets and a branching ductal network ending in exocrine glands. The endocrine compartment secretes hormones to regulate glucose homeostasis, while the exocrine pancreas produces enzymes that are involved in digestion. To accommodate both the endocrine and exocrine functions, the pancreas is organized into a complex network of ducts connected to terminally differentiated acini, with interspersed islets. During development, these three types of epithelial cells (ductal, acinar, endocrine) arise from a pool of multipotent pancreatic progenitors that are progressively specified into peripheral tip cells that will become acinar tissue and trunk cells that will give rise to ducts and islets^1^. Studies from model organisms have shown that surrounding mesenchymal cells provide physical and biochemical cues that are critical for proliferation and differentiation of epithelial progenitor cells^2–5^, as well as initiation of budding and branching morphogenesis of the ductal-acinar pancreas^6^.

Endocrine^7–9^ and exocrine^10–12^ organoids have been derived from human pluripotent stem cells (hPSCs). However, published methods generate only simple epithelial structures that do not contain the complete repertoire of functional endocrine, ductal, and exocrine tissues that are found in the native pancreas. One reason for the missing tissue complexity is likely the complicated nature of pancreas development, involving multiple different mesenchymal cell types that are required during the different stages of organogenesis. Previous studies have shown that including additional supporting cell types into organoid cultures results in elaborate tissue morphogenesis and the formation of complex liver and gastric tissues^13–15^. We therefore used a similar approach by including various mesenchymal cell types to better replicate human pancreas development, with the goal of generating pancreatic tissue organoids with a branching ductal network, functional exocrine glands, and endocrine islets.

### Identifying mesenchyme that promotes pancreas morphogenesis in vitro

In humans, the origin of the pancreatic mesenchyme and how it regulates morphogenesis are not known. However, evidence from chick and mouse models have suggested that different populations of mesodermally-derived cell types specify pre-pancreatic endoderm and the induction of the pancreatic bud^2,16,17^. These mesoderm populations include lateral plate and splanchnic mesenchyme, as well as vascular cell types like endothelial cells and pericytes. In fact, the majority of the pancreatic mesenchyme acquire a pericytic signature by E13.5 in mouse^18^. These studies support a potential role for splanchnic mesenchyme and pericyte-like mesenchymal progenitors in early pancreas development.

To generate complex pancreatic tissue organoids, we assessed the ability of different hPSC-derived mesenchymal populations to promote *in vitro* organogenesis. Given that all epithelial cell types in the pancreas derive from a common pancreatic progenitor that comes from the posterior foregut, we started with hPSC-derived dorsal posterior foregut endoderm (PFG)^8^. We tested two populations of hPSC-derived mesenchyme; splanchnic mesenchyme (SM)^19^ and a lateral plate mesoderm-derived population shown to have a high propensity for vascular and pericytic differentiation (LPM)^20,21^. We aggregated PFG cells with either day 4 SM to form splanchnic pancreas organoids (sPO) or day 5 LPM to form lateral plate/pericytic pancreas organoids (lpPO) (Fig. 1a). Endoderm and mesenchyme were aggregated in microwells at a 1:4 ratio, before being transferred to suspension culture with pancreatic medium containing retinoic acid and KGF, as well as BMP and Hedgehog inhibitors. At day 15, exogenous factors were removed, a step that was necessary for subsequent morphogenesis to occur. Although organoids recombined from both mesenchyme populations organized into a core of endoderm surrounded by mesenchymal cells, the epithelium in sPO rapidly formed large budding structures extending within the mesenchyme layer (Fig. 1b). In contrast, lpPO rearranged into a compact stratified epithelium similar to an embryonic pancreatic bud^22,23^, and this coincided with maintenance of strong expression of PDX1 and upregulation of the pancreatic transcription factor NKX6.1 (Fig. 1c,d). By day 25, 88±5% of the endodermal population expressed NKX6.1 whereas only 7±4% still expressed the foregut and stomach marker SOX2 (Fig. 1c,d). In sPO, SOX2 was abundantly expressed while the NKX6.1+ population accounted for only 19%±5% of the endoderm cells. These results suggest that LPM drives PFG fate towards the pancreatic lineage.

**Figure 1:**
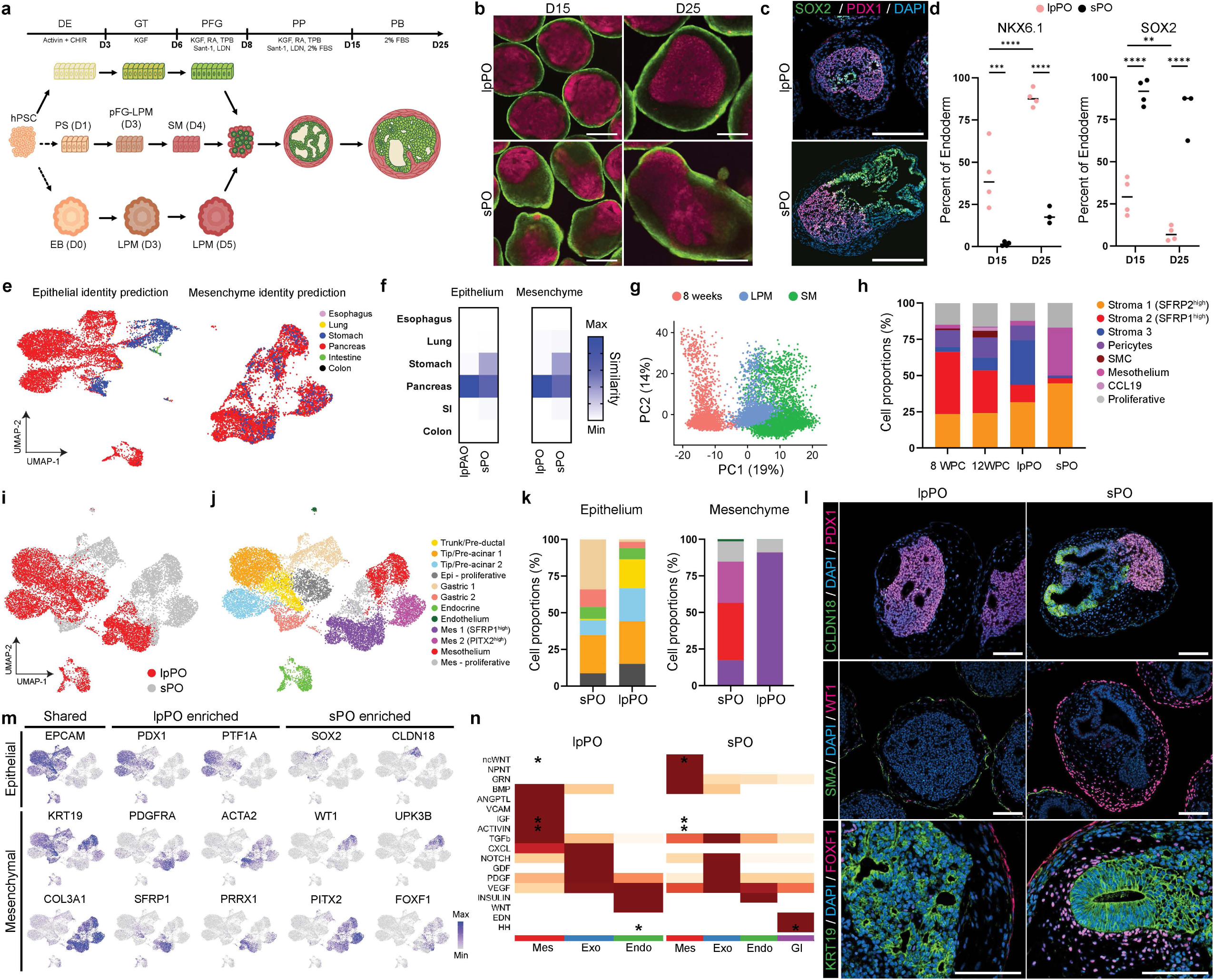
Mesenchyme identity biases pancreas morphogenesis. **a**, Schematic summary of the pancreatic organoid protocols using different mesenchyme populations. DE, definitive endoderm, GT, gut tube, PFG, posterior foregut, PP, pancreatic progenitors, PB, pancreatic bud, PS, primitive streak, LPM, lateral plate mesoderm, SM, splanchnic mesenchyme. **b**, Representative fluorescent images of the pancreatic organoids at days 15 and 25, with mesenchyme constitutively expressing GFP (green) and endoderm constitutively expressing mCherry (pink). Scale bars, 200 µm. **c**, Representative immunofluorescence staining of organoids at day 25. Scale bars, 200 µm. **d**, Intracellular flow cytometry analysis for NKX6.1 and SOX2 in the pancreatic organoids at day 15 and 25. Mean and standard deviation are shown for 4 independent differentiations. **p < 0.01, ***p < 0.001 and ****p < 0.0001, determined by two-way ANOVA with uncorrected Fisher’s LSD test. **e**, Uniform manifold approximation and projection (UMAP) for dimension reduction of scRNA-seq data showing identity prediction of the organoids against endodermal atlas^25,34^. **f**, Similarity score between organoids and different organs from the endodermal atlas^25,34^, obtained using a machine learning algorithm for identity prediction. **g**, Principal Component Analysis (PCA) of SM, LPM and fetal pancreatic mesenchyme from 8 wpc^25^. **h**, Bar graph showing the proportions of cells that are mapped to specific cell types from human fetal pancreatic mesenchyme, with annotations modified from *de la O et al.*^25^. **i-j**, UMAP of scRNA-seq datasets for lpPO and sPO at day 25, colored by experimental conditions (**i**) and identified clusters (**j**). **k**, Graph showing proportions of cells in each cluster for the epithelium or the mesenchyme. Colors correspond to cell clusters in **j**. **l**, Representative immunofluorescence staining of organoids at day 25. Scale bars, 200 µm. **m**, Feature plots showing expression levels of indicated genes. **n**, Predicted outgoing cell-cell crosstalk between major cell types in sPO and lpPO as determined by CellChat. Asterisks denote known pathway critical to pancreatic development.

From these findings, we predicted that pancreatic organoids generated with LPM (lpPO) would be more similar to native developing human fetal pancreas than those generated with splanchnic mesenchyme (sPO). To investigate this, we performed single-cell RNA-sequencing (scRNA-seq) on lpPO and sPO at day 25, and benchmarked these data to a scRNA-seq atlas of endodermal organs including pancreas and other surrounding organs (esophagus, lung, stomach, small intestine, colon). A machine learning identity prediction algorithm concluded that lpPO mapped almost exclusively to the pancreas (similarity score = 96%) (Fig. 1e,f). In contrast, sPO additionally contained a large population of gastric epithelial cells. Mesenchymal identity was also predicted to be highly pancreatic in the lpPO, whereas sPO mesenchyme additionally shared similarities with stomach mesenchyme. Consistent with this result, single-cell principal component analysis (Fig. 1g) and identity mapping (Fig. 1h) confirmed that the lpPO mesenchyme was more similar to human fetal pancreas compared to sPO mesenchyme.

Unbiased clustering revealed 7 epithelial clusters and 4 mesenchymal clusters distributed amongst the two different organoid datasets (Fig.1i,j). Notably, the gastric-like clusters (*CLDN18*, *SOX2*) were almost exclusively represented in the sPO, whereas lpPO was highly represented in clusters with pancreatic progenitor gene expression (*PDX1*, *PTF1A*, *CPA1*, *SOX9*) (Fig. 1k-m). Moreover, the lpPO mesenchyme recapitulated the cellular diversity seen in the fetal pancreas, with similar proportion of cells being annotated as pericytes, smooth muscle cells and stromal cells. Two mesenchymal clusters were exclusively present in sPO, with one of them characterized by high expression of mesothelial markers (*WT1*, *KRT19*, *UPK3B*) (Fig. 1k,m). SM also had higher FOXF1 expression at both RNA and protein level, correlating with increased expression of Hedgehog target genes in the developing gastric epithelium (Fig. 1l,m). The mesenchyme transcriptome was more homogeneous in lpPO (1 cluster, *PRXX1*^high^/*SFRP1*^high^), with peripheral cells expressing higher levels of smooth muscle actin (Fig. 1l-m).

To identify why LPM and SM have differential capacity to promote pancreatic fate, we analyzed the transcriptome of the two mesenchyme populations pre-and one week post-recombination, the stage when pancreatic fate is specified. Bulk RNA-sequencing (RNA-seq) analysis followed by principal component analysis revealed that most of the variance (61%) was due to different mesenchymal identity obtained from the two protocols (Extended Data Fig.1a). One of the main differences was in signaling pathway components that are critical for early stages of pancreas development (Extended Data Fig. 1b-e). Signaling components that promote pancreatic fate that are expressed by LPM include Activin and IGF. In contrast, SM expressed Hedgehog, a pathway known to promote gastrointestinal and repress pancreatic fate. Moreover, LPM showed a more posterior identity based on HOX transcription factor levels (Extended Data Fig. 1e), and had higher expression of many genes typically associated with pancreatic mesenchyme (*PRRX1*, *PDGFRB*, *EMILIN2* and *ISL1*)^24–26^. In contrast, SM displayed higher expression of genes associated with intestinal or stomach mesenchyme, such as *CRABP1*, *OSR1,* and *OSR2* (Extended Data Fig. 1d).

CellChat cell-cell communication inference^27^ supported the idea that bi-directional communication between epithelium and mesenchyme may be driving pancreas versus gastric cell fate (Fig. 1n). In lpPO, Activin and IGF signaling were significantly upregulated, in accordance with their critical role during pancreas fate determination and subsequent progenitor proliferation^2,28^. The mesenchyme source of Activin signaling in lpPO appears to be mural-like cells (*ACTA2*^high^/*PDGFRB*^high^), one of the main sources of Activin in the human fetal pancreas^25^. In contrast, sPO involves epithelial hedgehog signaling acting on mesenchyme, and mesenchyme acting as a source of non-canonical WNT ligands (*WNT5B*) for the epithelium (Fig. 1n, Extended Data Fig. 1f-j), consistent with known roles of these pathway in stomach development^29^. Moreover, sPO mesothelium expresses inhibitors of pancreas development, like the Activin antagonist follistatin (Extended Data Fig. 1j).

We have validated that this protocol successfully generates organoids across five cell lines tested (two embryonic hPSC and three induced hPSC lines) (Fig.2, Extended Data Fig. 2a-d). As with all protocols, the quality of pancreatic organoids was directly dependent on the quality of the two starting populations, especially LPM differentiation. Suboptimal differentiation is manifested by an increase in other gastrointestinal cell types, especially gastric epithelium, that forms a dominant lumen at the expense of pancreatic tissue. However, in these cases, pancreatic differentiation efficiency could be rescued by exogeneous addition of Activin A during the first week in pancreatic medium (Extended Data Fig. 2b-d), consistent with our cell-cell communication analysis (Fig. 1n, Extended Data Fig.1). Conversely, addition of WNT3a shifted organoids towards gastric fate, while addition of the WNT inhibitor C-59 favored pancreatic differentiation. However, WNT inhibition also dramatically reduced organoid size (data not shown), likely via similar mechanisms to the precocious differentiation coupled with reduced exocrine mass seen in Pdx1-Cre β-catenin floxed mice^30^. These results demonstrate that manipulation of Activin and WNT signaling can be used to push the PFG towards gastric or pancreatic organoid.

**Figure 2:**
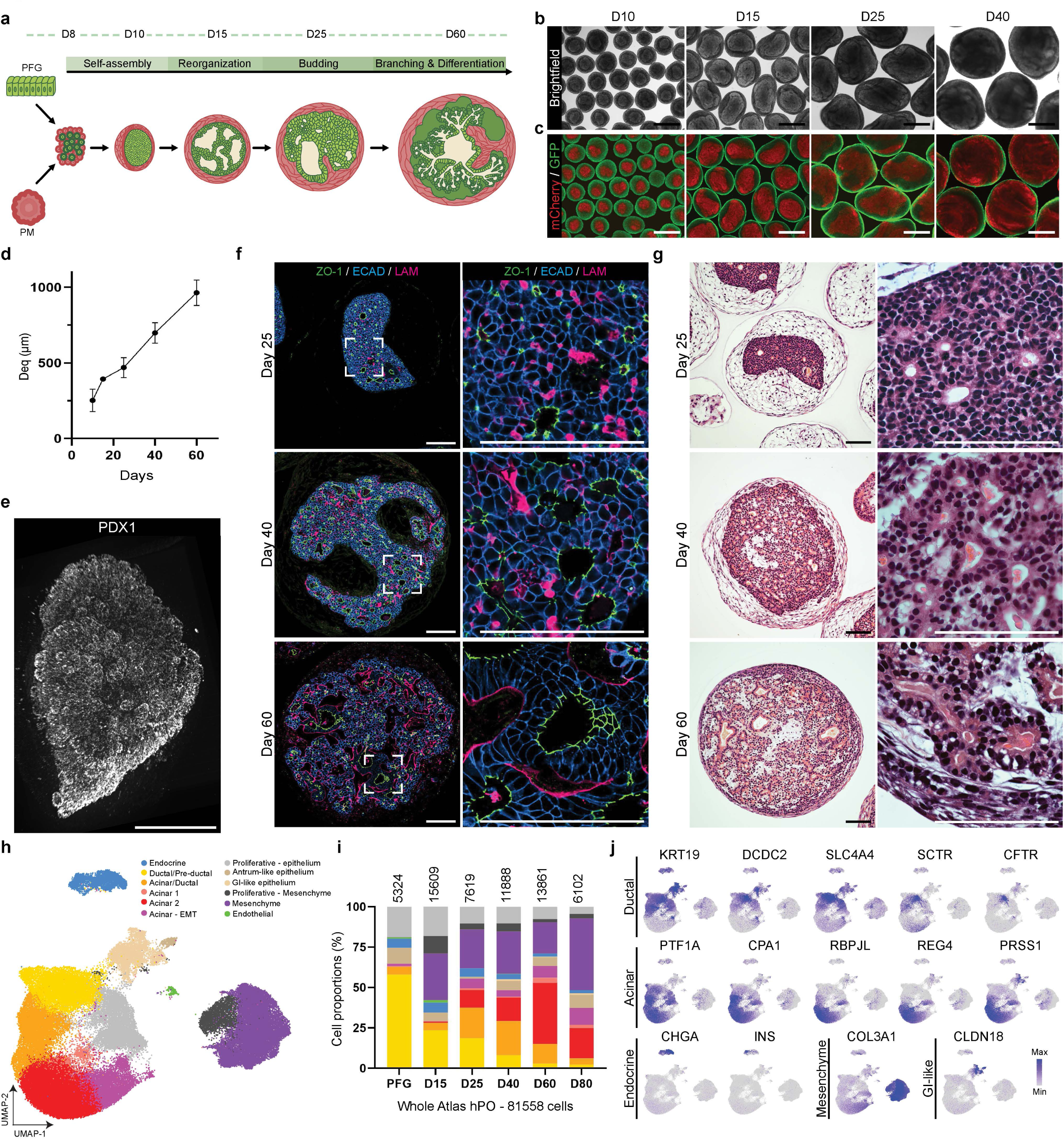
Pancreatic organoids undergo complex morphogenesis. **a**, Schematic summary of the pancreatic organoid (hPO) protocol. **b-c**, Representative brightfield (**b**) and fluorescence (**c**) images of the organoids at different timepoints. Scale bars, 400 µm. **d**, Graph showing hPO diameter in function of time in culture. The first point represents organoids at day 9 (one day after recombination). Mean and standard deviation are shown for 3 independent differentiations. **e**, 3D visualization of whole mount immunostaining of hPO at day 60. Scale bar, 400 µm. **f-g**, Representative images of immunostaining (**f**) and Hematoxylin and Eosin staining (**g**) of the hPOs at different timepoints, showing whole organoids (left) and magnified views (right). Scale bars, 100 µm. **h**, UMAP of scRNA-seq data from the whole hPO atlas incorporating 6 different timepoints, color-coded according to cell type annotations. **i**, Bar graph showing the cell proportions for each cell type throughout the differentiation. Colors correspond to cell clusters in **h**. **j**, Feature plots showing expression levels of indicated genes.

### Human pancreatic organoids undergo morphogenesis akin to developing human pancreas

When combined with LPM, PFG cells rapidly coalesce into a dense core of cells surrounded by mesenchyme before undergoing complex cellular rearrangement and morphogenesis (Fig. 2a). Establishment and maintenance of this core-mantle configuration was essential for successful initiation of a pancreatic bud-like program and later formation of pancreatic organoids (Fig. 2b,c). Similar to the developing pancreas, the pancreatic epithelium rapidly expanded relative to the mesenchyme, with the epithelial to mesenchymal cell ratio progressively increasing throughout organoid development (Extended Data Fig. 3a). One week after recombination of LPM and PFG endoderm, all exogenous factors were removed from the cultures, with all subsequent development of pancreatic tissues depending on intrinsic communication between the epithelium and mesenchyme. Flow cytometry analysis revealed that 90% of epithelial cells were pancreatic, with less than 10% of the cells expressing either SOX2 or CDX2 at day 25 (Extended Data Fig. 3b)^31^. PDX1 was maintained in 95±3% and 97±1% of the epithelial cells at day 15 and 25, respectively. Immunostaining analysis revealed colocalization of PDX1, SOX9, and NKX6.1 together with a high proportion of Ki67^+^ cells (Extended Data Fig. 3c,d), marking multipotent pancreatic progenitors reported to give rise to all epithelial cells of the developing pancreas^1^.

Human pancreatic organoids (hPO) grew and developed for over 2 months in culture, forming large branching structures that reached a millimeter in diameter (Fig. 2d,e). During this culture period, hPOs underwent a series of morphogenetic events that were strikingly similar to pancreas development *in vivo*. By day 25, hPOs displayed a stratified epithelium with multiple microlumens (Fig. 2f), like the embryonic pancreatic bud^23^. Over time, these microlumens became progressively contiguous, forming a network of connected epithelial branches, a process that mirrors the development of the ductal plexus in vivo^22,23^. By day 60, epithelial apical-basal polarity was well established, with ZO-1 on the apical side and laminin on the basal side. Hematoxylin and Eosin staining confirmed that hPOs stages were similar to the developing pancreas, with stratified epithelium transitioning to a cuboidal/ductal epithelium, and elongated ducts leading to polarized acinar-like cells with basally positioned nucleus organized around a lumen (Fig. 2g).

To comprehensively investigate if hPO physiologically mirrored human pancreas development, we generated a single-cell atlas of hPO development containing 80,289 cells derived from six different timepoints (PFG day 8 and hPO day 15, 25, 40, 60, 80) (Fig. 2h-j, Extended Data Fig. 3, Extended Data Fig. 4). The hPO atlas was then benchmarked to several single-cell datasets from the developing human fetal pancreas between 7-20 weeks gestational age^7,24–26^. The initial population of posterior foregut cells transitioned to committed pancreatic progenitors by D25 (Extended Data Fig. 3e-i). Pancreatic progenitors formed 3 clusters that were also found in the human early fetal pancreas^7^. One cluster expressed higher level of trunk cell (*SPP1*, *DCDC2*) and functional duct (*SLC4A4*, *CFTR*) markers, while the two other clusters correlated with tip cell identity and expressed the early exocrine markers *PTF1A* and *CPA1* (Extended Data Fig. 3h,i). At later stages (day 40, 60, 80) we observed ductal differentiation as marked by *CFTR*, *SCTR,* and *SLC4A4* and acinar differentiation as marked by expression of digestive enzymes such as *PRSS1* and *CPA1* (Fig. 2j, Extended Data Fig. 4a-e). The formation of endocrine cell was observed early at D15 as marked by NEUROG3 and Chromogranin A, with formation of cells positive for islet hormones like insulin and glucagon (Fig. 2j, Extended Data Fig. 4a,e). Notably, only 6.4% of the cells showed a non-pancreatic signature, based on expression of gastric markers (*CLDN18*, *SOX2*) and intestinal markers (*CDH17*, *CDX2*). Mesenchymal cells accounted for 24.6% of the cells and showed highest expression of *VIM*, *ACTA2*, *PDGFRA,* and *COL3A1*. From these analyses, we conclude that hPOs contain all the expected epithelial cell types found in the developing human pancreas and that their proportions broadly match what is expected from pancreatic organogenesis.

### Pancreatic organoids form a spatially organized ductal network terminating in functional acini

Analysis of ductal/exocrine development in hPOs revealed 14 clusters labelled as proliferating, ductal/progenitors, and acinar cells based on published scRNA-seq dataset (Fig. 3a-d)^7,24,25^. Pseudotime analysis suggested that pancreatic progenitors first form a progenitor population with shared ductal-acinar gene expression (*SLC4A4*^+^/*CPA1*^+^) that then separates into ductal and exocrine populations that undergo maturation over time. These results were confirmed by the inferred timecourse from the whole atlas (Fig. 3b-d). Based on transcripts for exocrine enzymes, we observed 6 populations of acinar cells that subclustered based on early to late developmental stages (clusters 3, 8, 0, 1, 10, 11). Early exocrine cells expressed genes like *SUSD4* and *CPA1,* whereas more mature exocrine markers like *PRSS1, RBPJL,* and *REG4* were expressed at later stages. We also observed that not all enzymes were expressed in the same cells, suggesting heterogeneity in enzyme expression within different acinar subtypes.

**Figure 3:**
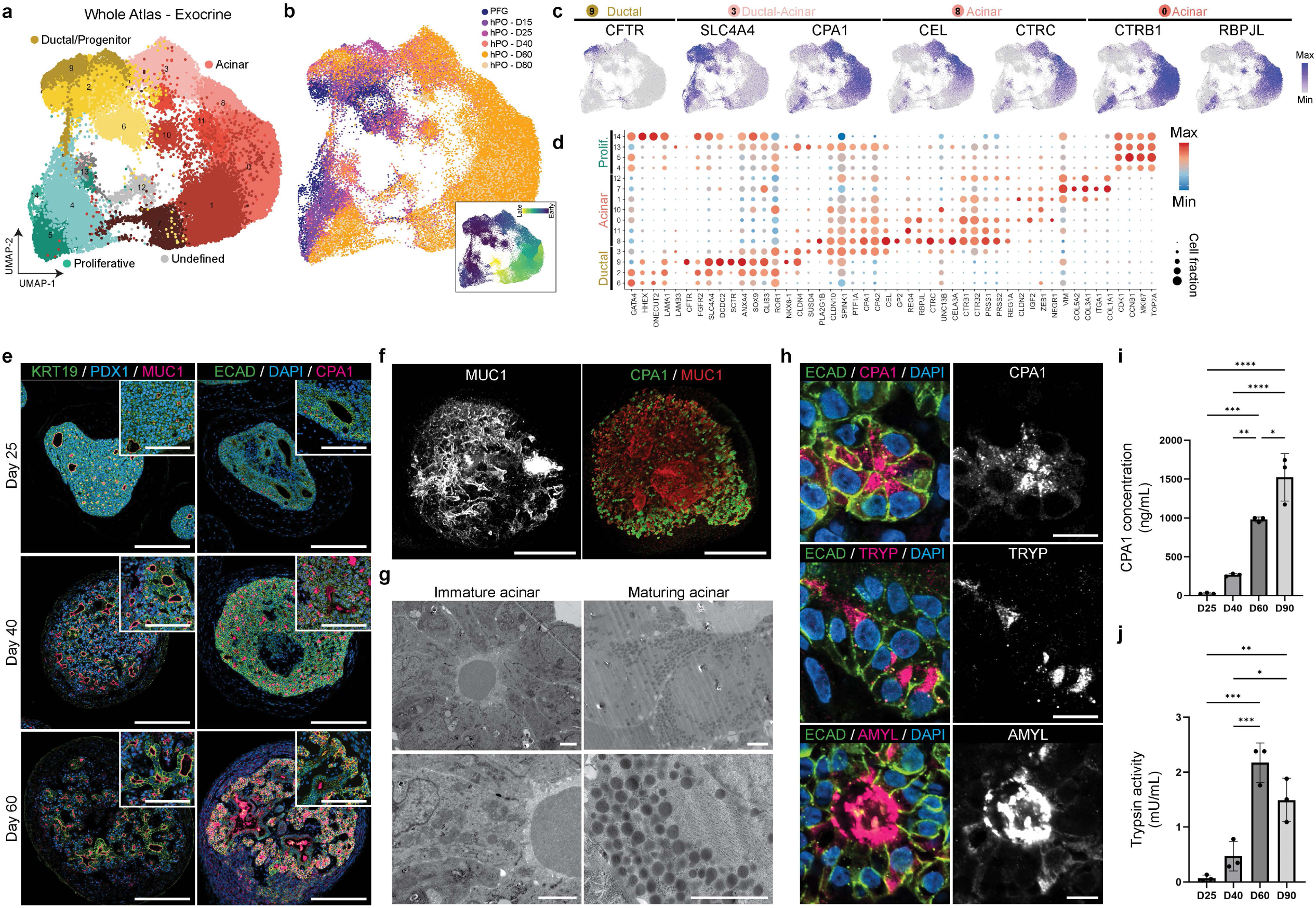
hPOs have spatially patterned ducts and functional acini. a-b, UMAP projections of subsetted exocrine cells from the hPO atlas, with cell identity annotated by broad cell types (**a**) or timepoint (**b**). Insert in **b** represents inferred timecourse from pseudotime analysis. **c**, Feature plots showing expression levels of indicated genes. **d**, Dot plot showing the expression of key marker genes across cell clusters. Dot sizes and colors indicate proportions of cells expressing the corresponding genes and their averaged scaled values of log-transformed expression, respectively. **e**, Representative immunostaining images of organoids at different timepoints showing mucin and enzyme expression. Scale bars, 200 µm. **f**, 3D visualization of whole mount immunostaining of hPO at day 60. Scale bars, 200 µm. **g**, Representative electron micrograph of day 60 organoids, showing immature acinar (left) and maturing acinar (right) cells with dense zymogen granules. Scale bars, 2 µm. **h**, Representative immunostaining images of acinar cells in day 60 organoids, showing merged pseudocolor (left) and grayscale for the selected enzyme (right). Scale bars, 10 µm. **i-j**, Graph quantifying secretion of enzymes into organoid lumen over time, showing CPA1 concentration (**i**) and trypsin activity (**j**). Mean and standard deviation are shown for 3 independent differentiations, with 10 organoids pooled from each differentiation. *p<0.01, **p < 0.01, ***p < 0.001, and ****p < 0.0001, determined by one-way ANOVA with Tukey’s multiple comparisons test.

The developing exocrine pancreas is spatially organized into a contiguous ductal network that terminates in exocrine clusters oriented for luminal secretion^23^. hPOs were similarly organized into an interconnected ductal network expressing KRT19 and apical secretion of MUC1 (Fig. 3e). While the exocrine protein CPA1 was initially expressed at low level in day 25 hPOs and was largely intracellular, we observed an increase in secreted CPA1 over time (Fig. 3e). Importantly, whole mount immunostaining confirmed *in vivo*-like spatial distribution of ductal and acinar tissue, with centrally localized larger ducts leading to smaller branching ducts terminating in acini that were predominantly located in the periphery of the organoids (Fig. 3f). Electron microscopy of day 60 hPOs showed that some acini have an immature phenotype, with basally located nuclei and light zymogen granules (Fig. 3g). In addition, multiple acinar cells had a more mature phenotype, with dense zymogen granules apically positioned near the lumen, and well-organized endoplasmic reticulum (Fig. 3g). Immunofluorescence staining for pancreatic enzymes like CPA1 also showed apical localization of zymogen granules in acini (Fig. 3h). Other pancreatic enzymes such as trypsin and amylase were also detected at later stages of hPO development, similar to their expression during human fetal development, where trypsinogen is initially detected at 14-16 weeks post-conception (wpc) and amylase is expressed at low levels until birth^32,33^. Analysis of secretions from the organoids also demonstrated an increase in luminal CPA1 and trypsin levels during organoid development (Fig. 3i,j). Together, these data demonstrate that hPOs contain a complex ductal network that terminate in spatially organized and functional acini.

### Mapping the stages of hPO development onto human fetal pancreas development

Given that many organoid systems have off-target cell types, we performed an unbiased comparison of hPO against two different fetal atlases of human endodermal organs^24,25,34^ using a machine learning algorithm (Extended Data Fig. 5a-d). Consistent with our flow cytometry analysis (Extended Data Fig. 3b), the vast majority of cells were classified as pancreatic, with a small proportion mapping closer to the fetal stomach (Extended Data Fig. 5e-l). Similarly, the hPO mesenchyme mapped onto fetal pancreatic mesenchyme, with a low similarity score to stomach (Extended Data Fig. 5f,j), consistent with the proximity between the developing stomach and pancreas mesenchyme.

Next, we projected the hPO data onto the fetal pancreas atlas^25^ in attempts to identify the relative cell proportion (Fig. 4a,b) and .developmental stages (Fig. 4c,d) of the epithelium in hPOs. At stages where PFG cells transition into pre-acinar progenitors (day 15-25), hPOs were most similar to 8 wpc embryos, the earliest timepoint available in the fetal atlas. As hPO ductal and acinar differentiation proceeded (days 40-60), organoids were more similar to 12 wpc human embryos (Fig. 4c,d). By day 80, hPOs had a population of cells that mapped onto samples from 19-20 wpc human embryos. A similar analysis of the mesenchymal cells in hPOs revealed 5 clusters, including a proliferative cell cluster and 4 different mesenchymal populations (Extended Data Fig. 6a-c). Mes 1 represents a relatively immature population, mapping most closely to pericytes when compared to the human fetal atlas (Fig. 4e,f, Extended Data Fig. 6d). Mes 2, 3, and 4 were more similar to the three stromal populations found in the fetal atlas (Extended Data Fig. 6d) and broadly expressed *SFRP1*, *PRRX1*, *POSTN*, *COL3A1*, and *MGP* (Extended Data Fig. 6e,f). hPO mesenchyme also underwent maturation over time, with early cells mapping onto 8 wpc human embryos whereas later organoids mapped onto 12 wpc samples (Fig.4g,h). Overall, these data suggest that hPO epithelium and mesenchyme both continued to mature in culture over time, and that mesenchyme had similar complexity than endogenous mesenchyme (Fig. 4e,f).

**Figure 4:**
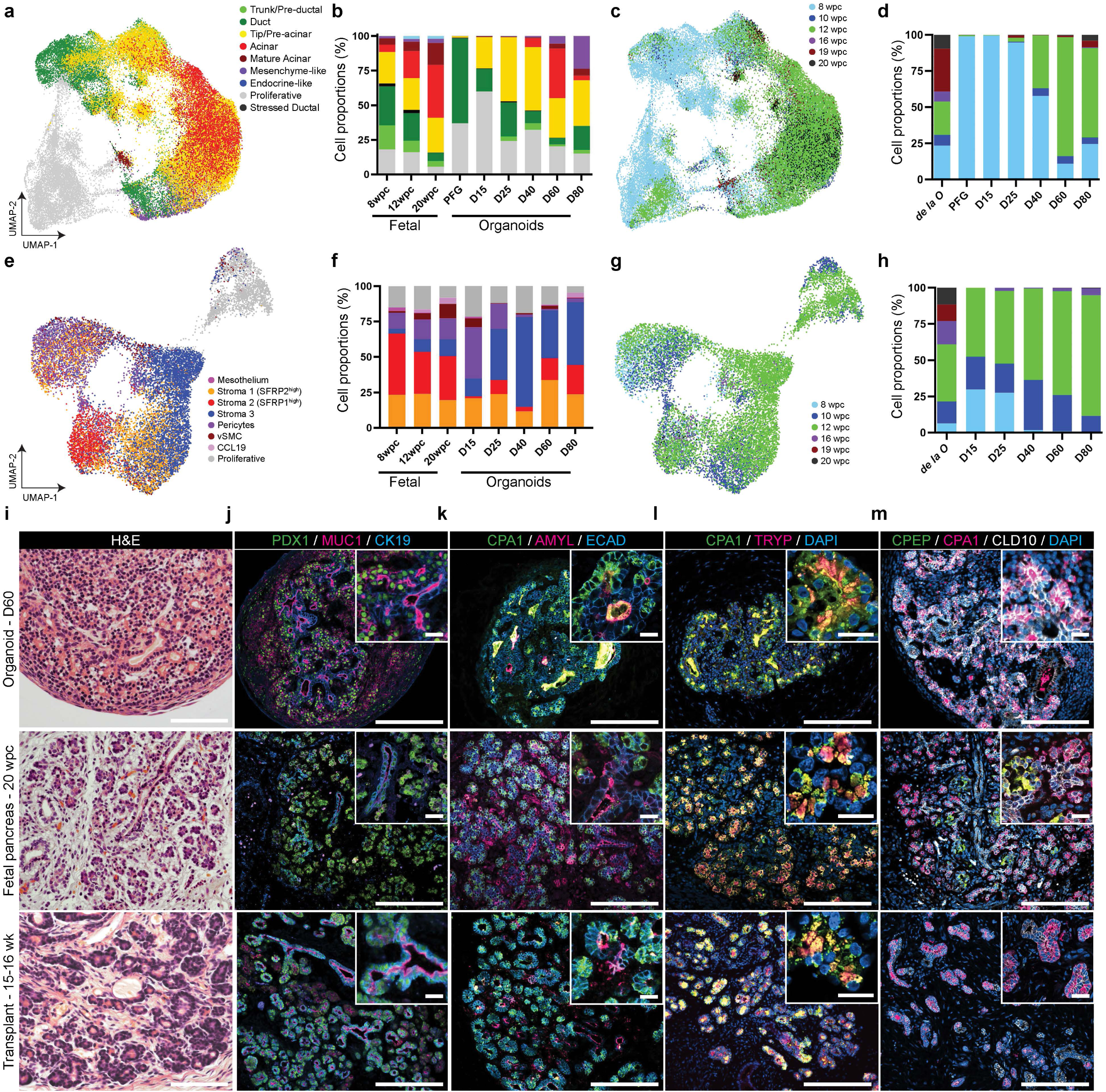
hPOs recapitulate human development. **a**, UMAP of scRNA-seq data showing identity prediction of organoid’s exocrine lineage against a fetal pancreas atlas^25^, color-coded based on annotations of cell types in the fetal pancreas. **b**, Bar graph showing cell proportions in exocrine tissue for each organoid developmental timepoint, color-coded based on cell type annotations used in **a**. **c**, UMAP of scRNA-seq data showing age prediction of the organoid’s exocrine lineage against the human fetal pancreas atlas from *de la O, et al.*, color-coded according to the corresponding age in the fetal atlas. **d**, Bar graph showing cell proportions in exocrine tissue for each organoid developmental timepoint, color-coded based on age annotations used in **c**. **e,** UMAP of scRNA-seq data showing identity prediction of organoid’s mesenchymal lineage against a fetal pancreas atlas, color-coded based on annotations of cell types in the fetal pancreas. **f**, Bar graph showing cell proportions in mesenchymal tissue for each organoid developmental timepoint, color-coded based on cell type annotations used in **e**. **g**, UMAP of scRNA-seq data showing age prediction of the organoid’s mesenchymal lineage against the human fetal pancreas atlas from *de la O, et al.*, color-coded based on the corresponding age in the fetal atlas. **h**, Bar graph showing cell proportions in mesenchymal tissue for each organoid developmental timepoint, color-coded based on age annotations used in **g**. **i**, Representative Hematoxylin and Eosin staining images of the hPOs at day 60 (top), fetal pancreas at 20 wpc (middle) and transplanted hPO (bottom). Scale bars, 100 µm. **j-m**, Representative immunostaining images of hPOs (top), fetal pancreas (middle) and transplanted hPOs (bottom). Inserts show a magnified view to highlight protein distributions at the microscopic scale. Scale bars, 200 µm (main) and 20 µm (insert).

Comparison of late stage hPOs to 20 wpc pancreatic fetal sections revealed some similarities (Fig. 4i-m), with hPOs containing cuboidal ducts and acini expressing exocrine markers. However, the acinar and ductal structures in hPOs were more compact and less defined than in 20 wpc pancreatic sections. Our transcriptomic data of day 60 hPOs places them between 12 and 16 wpc, and hPOs do not continue to mature substantially past 90 days *in vitro*. To enable continued maturation, we transplanted day 25 organoids under the kidney capsule of immunocompromised (NSG) mice and grew them for an additional 3 months (total 15-16 weeks of age). Previous studies have shown that growth of PSC-derived intestinal and gastric organoids *in vivo* facilitates continued growth and maturation^13,35^. Transplanted hPOs had strikingly similar architecture and protein distribution of CPA1, trypsin, and amylase, demonstrating that hPO continue to develop and mature when provided with an adequate environment (Fig. 4j-l). Overall, these data show that hPOs develop and mature into tissue that is akin to early second trimester human pancreas.

### Endocrinogenesis forms islet-like clusters within the organoids

Although scattered endocrine cells were present in hPOs, they failed to form islet-like endocrine clusters (Fig. 4m, Figure 5a,b). Pancreatic mesenchyme has been shown to bias multipotent pancreatic progenitors towards an exocrine fate^36^, suggesting that extrinsic endocrine-inducing factors may be needed to promote endocrine and islet development in hPOs. Addition of pancreatic progenitor inducing media (PP medium, Fig. 1a) during the first week of hPO culture was required to promote gene expression programs characteristic of endocrine progenitors (*SUSD2*, *NEUROG3*, *INSM1*) and islet hormone expression (*SST*, *GHRL*) (Extended Data Fig. 7a-c). That said, endocrine cells did not form islet like clusters and were not robustly maintained at later timepoints (Fig. 5a,b), suggesting that signals that promote endocrine islet development are not produced in our organoids. We therefore modulated signaling pathways known to promote endocrine differentiation from pancreatic progenitors^8,9^. Inhibition of Notch (X), BMP (L) and hedgehog (S) signaling, and activation of thyroid hormone signaling (T) for 10 days promoted differentiation of endocrine progenitors expressing CHGA and NEUROG3 in day 25 organoids, later resulting in the formation of islet-like clusters that expressed the pancreatic hormone glucagon and C-peptide (Fig. 5a,b).

**Figure 5:**
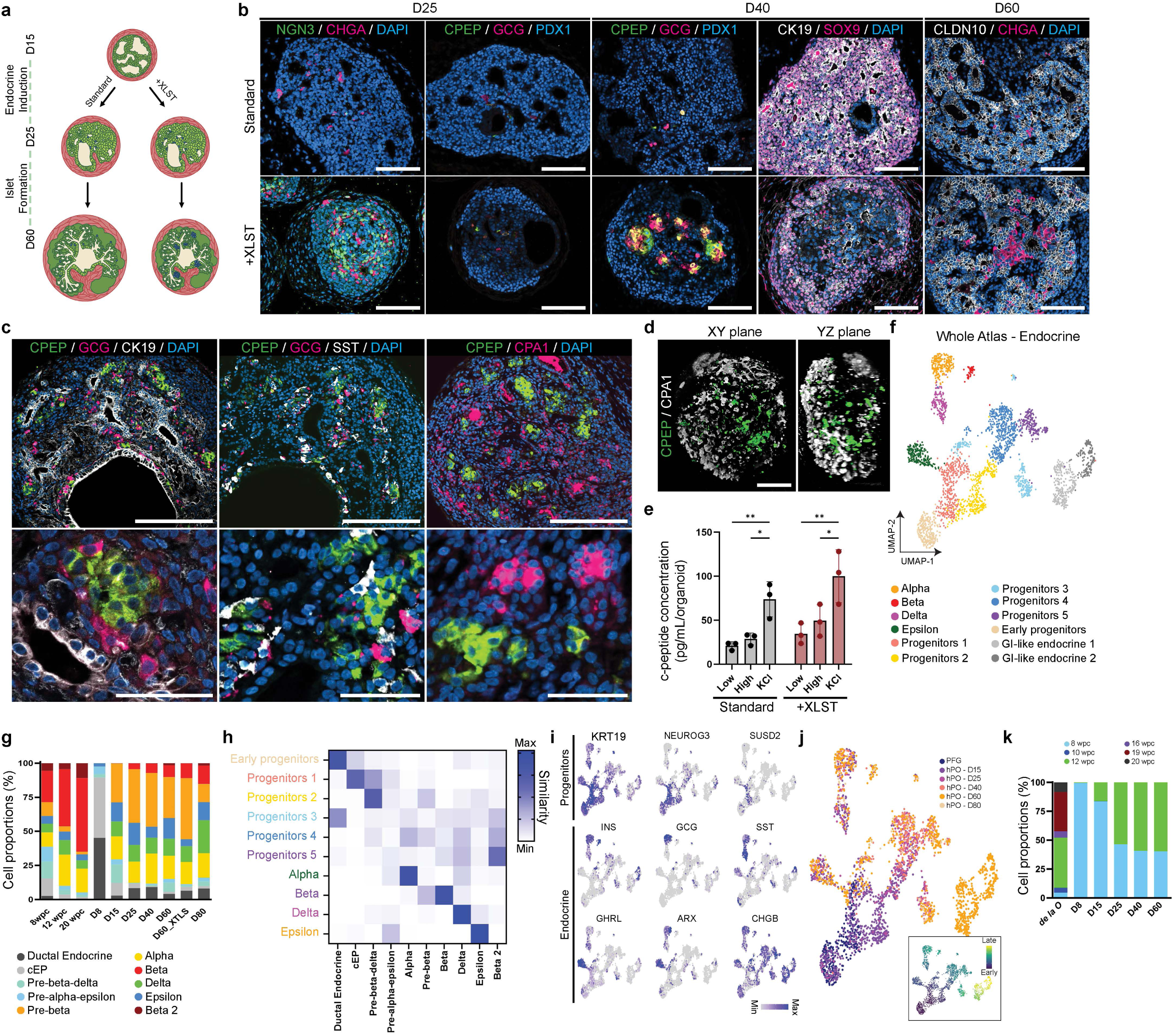
Islet-like clusters form within hPOs. **a**, Schematic summary of the endocrine differentiation protocol. **b**, Representative immunofluorescence images of the organoids at different timepoints based on the standard protocol or addition of the endocrine differentiation cocktail *XLST*. Scale bars, 100 µm. **c**, Representative immunofluorescence images of endocrine-induced day 60 organoids, showing low (top) and high (bottom) magnifications. Scale bars, 200 µm and 50 µm for low and high magnification, respectively. **d**, 3D visualization of whole mount immunostaining of endocrine-induced hPO at day 60. Scale bar, 200 µm. **e**, Graph showing KCl-and glucose-stimulated release of c-peptide, a proxy for insulin, from the organoids. **f**, UMAP projections of subsetted endocrine cells from the hPO atlas, color-coded according to cell type annotations. **g**, Bar graph showing identity prediction of endocrine cell types against fetal endocrine pancreas atlas, color-coded based on annotations of cell types present in the fetal pancreas^25^. **h**, Similarity score between the endocrine cluster in the hPO atlas (y axis) and endocrine cells from the fetal pancreas atlas (x axis)^25^, obtained using a machine learning algorithm for identity prediction. Cell types are color-coded based on populations present in the whole endocrine atlas (**f**). **i**, Feature plots showing expression levels of indicated genes. **j**, UMAP projections of endocrine cells from the whole atlas, with cell identity annotated by timepoint. Insert represents inferred timecourse from pseudotime analysis. **k**, Bar graph showing predicted proportions of endocrine cells within the organoids that match specific fetal pancreas timepoints^25^.

Single-cell transcriptome analysis of day 60 organoids treated with a pulse of XLST confirmed increased endocrine differentiation, with increased expression of both progenitor cell (*CHGA*, *NEUROG3*, *SUSD2*) and more mature islet cell markers (*INS*, *GCG*, *SST*, *MAFB*, *CHGB*) (Extended Data Fig. 7d-g). Interestingly, XLST pulse was sufficient to maintain large number of CHGA+ cells throughout organoid development until at least day 60 (Fig. 5b). At that stage, islet clusters could be found adjacent to the ducts, with cells displaying expression of c-peptide, glucagon, and somatostatin (Fig. 5c). Many of these cells were mono-hormonal, a feature consistent with more mature islet cells. Whole mount immunostaining showed that most islet clusters were located in the center of the organoids, in contrast to the CPA1+ acinar cells located at the periphery (Fig. 5d), similar to the spatial organization of endocrine and exocrine tissue in vivo^37–39^. Pancreatic organoids secreted increased insulin in response to KCl depolarization or step changes in glucose concentration, although it did not reach statistical significance between low and high glucose conditions (Fig. 5e). Nevertheless, this suggests that endocrinogenesis can be recapitulated within the organoid, and that it results in islet clusters that show functional maturation.

Reclustering the endocrine cells from the hPO atlas revealed 12 endocrine clusters that were annotated based on expression in human fetal pancreas atlas (Fig. 5f)^24–26^. HPOs contained all four major islet cell types normally present in the dorsal pancreas: alpha (*GCG*, *IRX2*), beta (*INS*, *MAFA*), delta (*SST*, *HHEX*), and epsilon (*GHRL*, *ARX*) cells. Using similarity scoring against human fetal pancreatic endocrine cells, we confirmed the identity of each of the hPO cell types, including alpha, beta, delta, epsilon, and progenitor cells (Fig. 5g,h). The remaining endocrine clusters mapped to endocrine progenitors, expressing ductal (*KRT19*) and endocrine progenitor (*NEUROG3*, *SOX4*, *PAX4*, *SUSD2*) markers (Fig. 5h,i). To investigate if endocrine differentiation in hPOs was similar to human fetal pancreas, we performed pseudotime analysis with Monocle, which showed bifurcation from a common endocrine progenitor (cEP) to pre-specified progenitor populations that progress towards the different islet cell types (Fig. 5j). It also confirmed a decrease in proportions of progenitors (ductal endocrine, cEP, Pre-beta) and an increase in proportions of islet cells that mimicked the relative changes in the fetal pancreas (Fig. 5g). Projection of the endocrine population onto the fetal endocrine atlas confirmed that endocrine progenitors mapped to early progenitors (8 wpc), whereas differentiated islet cells were most similar to 12 wpc (Fig. 5k). We conclude that concurrent exocrine morphogenesis and islet formation can be recapitulated within pancreatic organoids via conserved developmental principles.

## Discussion

Over the past decade, numerous organoid models of pancreatic ductal, exocrine, or endocrine have been generated from hPSCs^7–12,40^. Each of these consists of simple epithelial structures that do not capture the cellular and architectural complexity of the native pancreas. Here, we have established and extensively benchmarked a new pancreatic organoid that contains the epithelial complexity and functionality of the developing human pancreas. Although previous studies have combined mesenchymal and hPSC-derived endodermal cells in other organs like the liver, stomach or bladder^13,15,41^, our progenitor assembly approach reveals the importance of recapitulating time and organ specific epithelial-mesenchymal interactions to unlock the self-organization potential of pancreatic progenitors to form a branching ductal-acinar network. Overall, hPOs represent the first model to contain spatially patterned ductal, exocrine, endocrine, and mesenchymal tissue in a physiologically relevant way, opening new possibilities for testing cellular interactions during pancreatic development and diseases. In addition, hPOs represent a new platform to study a broad array of pancreatic diseases that impact ductal, exocrine, and endocrine components.

## Methods

### hPSCs Lines & Maintenance

Human embryonic stem cells WA01 (H1, RRID: CVCL_9771) and WA09 (H9, RRID:CVCL_9773) were obtained from WiCell. Human induced pluripotent stem cells were obtained from either Coriell Institute for Medical Research (WTC CDH5-eGFP, AICS-0126-041) or from the CCHMC Pluripotent Stem Cell Facility (iPSC72.3 -RRID:CVCL_A1BW, iPSC263.10, iPSC1096.8). hPSC were maintained in mTeSR1 (StemCell Technologies, 85850) and cultured on Nunc™ tissue culture treated 6-well plates (ThermoFisher Scientific, 140675) coated with hESC-qualified Matrigel (Corning, 354277). hPSCs were passaged every 3-4 days using 3-5 min treatment in PBS without Mg2+ and Ca2+ (ThermoFisher Scientific, 10010023) supplemented with 0.5 mM EDTA (Invitrogen, 15575020), followed by colony detachment using a cell scraper. All hPSC were confirmed karyotypic normal and were expanded into a working cell bank frozen in mFreSR™ (StemCell Technologies, 05955) and stored in liquid nitrogen. hPSC were routinely confirmed negative for mycoplasma using EZ-PCR Mycoplasma Test kit (Sartorius, 2070020).

### Generation of posterior foregut cells

When reaching 70-80% confluency, hPSC colonies were rinsed with PBS followed by incubation with TrypLE Express (ThermoFisher Scientific, 12604021) for 3–4 min at 37 °C. Colonies were dissociated into single cells, collected, mixed with mTeSR medium, and spun at 1,000 rpm for 3 min. The pellet was resuspended in mTeSR medium supplemented with ROCK inhibitor Y-27632 (10 mM, Tocris, 1254) and seeded at a concentration of 1.5 × 10^5^ cells/cm^2^ on plates coated with 1:30 diluted Matrigel Growth Factor Reduced (Corning, 356231). Medium was changed after 24 hours without ROCK inhibitor. Plating density was adjusted for different cell lines to achieve 100% confluency after 48 hours of culture. The differentiation was carried out following previously established protocols with minor modifications^8,40^. Briefly, cells were differentiated towards definitive endoderm using endoderm basal medium supplemented with 100 ng/mL Activin A (BioTechne, 338-AC) and 3 µM CHIR99021 (CHIR, Tocris, 4423), and towards primitive gut tube using 50 ng/mL KGF (R&D Systems, 251-KG-CF) and 0.25 mM ascorbic acid (Sigma-Aldrich, A4544). Differentiation towards posterior foregut was achieved with pancreatic basal medium supplemented with 50 ng/mL KGF, 0.25 µM SANT-1 (Sigma-Aldrich, S4572), 1 µM retinoic acid (Sigma-Aldrich, R2625), 200 nM LDN193189 (ReproCell, 04-0074), 200 nM TPPB (Tocris, 5343). Endoderm basal medium consists of MCDB 131 medium (ThermoFisher Scientific, 10372019) further supplemented with 1.5 g/L sodium bicarbonate (Sigma-Aldrich, S6297), Glutamax (1X), 10 mM glucose (Sigma-Aldrich, G7528) and 0.5% fatty acid free Bovin Serum Albumin (BSA, Sigma-Aldrich, A7030). Pancreatic basal medium consists of MCDB 131 medium supplemented with 2.5 g/L sodium bicarbonate, Glutamax (1X), 15 mM glucose (Sigma-Aldrich, G7528), 2% fatty acid free Bovin Serum Albumin (BSA, Sigma-Aldrich, A7030), 0.25 mM ascorbic acid and ITS-X (1X, ThermoFisher Scientific, 51500056).

### Generation of splanchnic mesenchyme

The directed differentiation of hPSCs into splanchnic mesenchyme has been previously described^19^. Briefly, hPSCs were exposed to Activin A (30 ng/ml), BMP4 (40 ng/ml, R&D Systems, 314-BP), CHIR99021 (6 µM), FGF2 (20 ng/ml, R&D Systems, 233-FB), and PIK90 (100 nM, Tocris, 7902) for 24 hours. A basal media composed of Advanced DMEM/F12 (ThermoFisher Scientific, 12634028) supplemented with B-27 supplement without vitamin A (1X, Thermo Fischer Scientific, 12597010), N2 supplement (1X, ThermoFisher Scientific, 17502048), HEPES (15 mM, ThermoFisher Scientific, 15630080), Glutamax (1X, ThermoFisher Scientific, 35050061), and Penicillin-Streptomycin (100 U/ml, ThermoFisher Scientifics, 15140122) was used for this and all subsequent differentiation steps. Cells were then exposed to A8301 (1 µM, Tocris, 2939), BMP4 (30 ng/ml), retinoic acid (2 µM) and C59 (1 µM, Cellagen Technology, C76412s) for 24 hours. For splanchnic mesoderm generation, cells were cultured in A8301 (1 µM), BMP4 (30 ng/ml), C59 (1 µM), FGF2 (20 ng/ml), and RA (2 µM, Sigma-Aldrich) from Day 2 to Day 4.

### Generation of lateral plate mesenchyme

LPM was generated using a previously published protocol with minor modifications^20^. In brief, cells were washed with PBS and subsequently incubated with TrypLE for 4 min at room temperature. Cells were collected, span down at 1000 rpm for 3 min and resuspended in Aggregation media consisting of KnockOut DMEM/F12 (ThermoFisher Scientific, 12660012), 20% KnockOut Serum Replacement (ThermoFisher Scientific, 10828028), GlutaMAX (1X), NEAA (1X, ThermoFisher Scientific, 11140050), 55 µM β-mercaptoethanol (ThermFisher Scientific, 31350010), 100 U/ml Penicillin-Streptomycin, 50 mM Y-27632. Cells were seeded into Aggrewell™ 400 (StemCell Technologies, 34415) at 1000-1500 cells/aggregates. The next day (day 0), aggregates were collected and let to sediment by gravity, before being resuspended in mesoderm induction medium and transferred to suspension culture in ultra-low attachment plate (Corning, 3471) placed on an orbital shaker at 100 rpm. Mesoderm induction medium was composed of N2B27 media supplemented with 12 µM CHIR and 30 ng/ml BMP4. N2B27 media consists of Neurobasal (ThermoFisher Scientific, 21103049) and DMEM/F12 (ThermoFisher Scientific, 11330031) mixed in 1:1 ratio, supplemented with B27 supplement with vitamin A (1X, ThermoFisher Scientific, 17504044), N2 supplement (1X), GlutaMAX (1X), 55 mM b-mercaptoethanol, and 100 U/ml Penicillin-Streptomycin. After 3 days of culture (day 3), media was exchanged with N2B27 media supplemented with 2 µm Forskolin (StemCell Technologies, 1000249) and 100 ng/ml VEGF-A (R&D Systems, 293-VE). Mesoderm aggregates were cultured for 2 more days before being ready for recombination (day 5).

### Pancreatic organoid formation and culture

Posterior foregut cells (PFG, day 8), splanchnic mesenchyme (SM, day 4) and lateral plate mesoderm (LPM, day 5) were dissociated into single cells using TrypLE Express for 4 min (PFG), 10 min (LPM) or 10-12 min (SM) at 37 °C. For LPM, mesenchymal aggregates were rapidly fluxed through a 1 mL pipet tip after 5 min of incubation. Enzyme inhibition was achieved by combining the cells and TrypLE with basal pancreatic medium in a 1:2 ratio. Cells were spun down at 300 g for 3 min and resuspended in pancreatic medium supplemented with 50 ng/mL KGF, 0.25 µM SANT-1, 1 µM retinoic acid, 200 nM LDN193189, 200 nM TPPB, 2% Fetal Bovin Serum (FBS, Cytiva, SH3007002), and 20 mM ROCK inhibitor Y-27632. Cells were then seeded in Aggrewell™ 800 (StemCell Technologies, 34811) with an endoderm-mesenchyme ratio of 1:4 (6000 PFG with 24000 mesenchyme in each microwell). The next day, organoids were transferred to suspension culture in ultra-low attachment plate (1 Aggrewell well into 1 ULA well) and placed on an orbital shaker at 100 rpm. The pancreatic medium without ROCK inhibitor was changed everyday for 1 week (day 15). At day 15, pancreatic medium was replaced with medium composed of Advanced DMEM/F12, B-27 supplement without vitamin A (1X), N2 supplement (1X), HEPES (15 mM), Glutamax (1X), Penicillin-Streptomycin (100 U/ml), and 2% FBS. For endocrine induction between day 15 and 25, this medium was supplemented with “XLST” cocktail composed of 100 nM Gamma secretase inhibitor XX (EMD Millipore, 565790), 200 nM LDN193189, 0.25 µM SANT-1 and 1 µM T3 (Tocris, 6666). In both cases, medium was changed every other day, and organoids were diluted in more wells if the medium turned strongly yellow during long-term culture.

### Flow cytometry

Organoids were washed in PBS and dissociated using TrypLE Express for 10-20 min at 37 °C, with forceful pipetting every 4-5 min. After enzyme inhibition with basal medium, cells were spun down for 3 min at 300 g and resuspended in PBS supplemented with 2% FBS. Cells were incubated for 20 minutes with fixable viability dye (L34992, ThermoFisher Scientific) and washed once before being fixed and permeabilized using the Cytofix/Cytoperm kit (BD Biosciences, 554714) for 12 min. After washing, cells were resuspended in Perm/Wash buffer (BD Biosciences, 554714) and incubated with conjugated antibodies for 30 minutes: PDX1-PE (BD Biosciences, 562161) NKX6.1-AF647 (BD Biosciences, 563338), CDX2-PE (BD Biosciences, 563428), SOX2-V450 (BD Biosciences, 561610), CD326-FITC (BD Biosciences, 347197), AlexaFluor® 647 mouse IgG1 K Isotype Control (BD Biosciences, 557732), PE mouse IgG1 K Isotype Control (BD Biosciences, 554680). They were then washed with Perm/Wash buffer, resuspend in PBS with 2% FBS, and passed through a 40 µm strainer before analysis. The experiments were run on an Aurora spectral analyzer (Cytek) and analyzed with FlowJo v.10 (BD Biosciences). For the gating strategy, debris and doublets were removed from analysis using FSC/SSC gating, and dead cells were removed by gating the fluorescent cells having incorporated the fixable viability dye. Isotype controls and unstained controls were used for the gating.

### Tissue processing and immunocytochemistry

For paraffin section, organoids were collected, washed with PBS and fixed overnight at 4°C using 4% paraformaldehyde (Alfa Aesar, J19943-K2). Small organoids were embedded in Histogel (Richard-Allan Scientific, HG4000012) to facilitate embedding of multiple organoids per conditions. Tissue was paraffin processed and embedded by the Integrative Research Pathology Facility at CCHMC. Tissue was deparaffinized in xylene (Fisher Scientific, HC700), rehydrated in progressive ethanol-water solution, and antigen retrieved with citrate buffer (Sigma-Aldrich, C9999) at high pressure (commercial pressure cooker). Tissues were serially sectioned at a thickness of 7-8 mm onto Superfrost Plus glass slides (Fisher Scientific, 1255015). Routine Hematoxylin & Eosin (H&E) staining was performed by the Research Pathology Core at CCHMC. For cryogenic sections, organoids were fixed for 1-2 hours at 4 °C, washed with PBS and resuspended in 30% sucrose solution in PBS overnight. Tissues were then embedded in O.C.T. compound (Sakura Finetek, 4583) and serially sectioned at a thickness of 7-8 mm onto Superfrost Plus glass slides.

For staining, slides were permeabilized and blocked in 5% normal donkey serum (Jackson ImmunoResearch, 017000121) in PBST (PBS with 0.5% Triton-X100 (Sigma-Aldrich, T8787)) for 1 hour. Slides were then incubated in primary antibody at 4 °C overnight. Primary antibodies used in this study were: goat anti-PDX1 (1:200, R&D Systems, AF2419), mouse anti-NKX6.1 (1:100, DSHB, F55A10), rabbit anti-SOX9 (1:200, EMD Millipore, AB5535), sheep anti-neurogenin3 (1:200, R&D Systems, AF3444), rat anti-C-peptide (1:200, DHSB, GNID4), rabbit anti-glucagon (1:200, Cell Marque, 259R15), mouse anti-chromogranin A (1:200, DHSB, CPTC-CHGA-1), mouse anti-cytokeratin 19 (1:200 for paraffin section, ThermoFisher Scientific, MA515884), rabbit anti-mucin 1 (1:200, Sigma-Aldrich, HPA008855), mouse anti-CPA1 (1:200, OriGene Technologies, TA500053), rabbit anti-GATA4 (1:200, Santa Cruz Biotechnology, sc9053), rabbit anti-trypsin (1:200, ThermoFisher Scientific, PA5106876), rabbit anti-amylase (1:200, Sigma-Aldrich, HPA045394), mouse anti-CDH17 (1:500, Sigma-Aldrich, SAB1403654), rat anti-SOX2 (1:200, R&D Systems, AF2018), rabbit anti-CDX2 (1:200, Cell Marque, 235R16), rabbit anti-claudin 18 (1:200, Sigma-Aldrich, HPA018446), rabbit anti-KI67 (1:200, Cell Marque, 275R14), goat anti-CDH1 (1:400, R&D Systems, AF648), mouse anti-ZO1 (1:200, ThermoFischer Scientific, 339100), rabbit anti-laminin (1:200, Abcam, ab11575), rabbit anti-collagen I (1:200, Abcam, ab34710), rabbit anti-WT1 (1:100, Abcam, ab89901), mouse anti-SMA (1:200, Sigma-Aldrich, A5228), mouse anti-vimentin (1:200, Santa Cruz Biotechnology, sc373717), goat anti-FOXF1 (1:200, R&D Systems, AF4798). Secondary antibodies used to visualize proteins included donkey anti-mouse Alexa Fluor 488, 568 and 647 (1:500; ThermoFisher Scientific, A21202, A10037, A31571), donkey anti-goat Alexa Fluor 647 (1:500; Thermo Scientific, A21447), donkey anti-rabbit Alexa-Fluor 568 or 647 (1:500; Thermo Scientific, A10042, A31573), donkey anti-rat 488 (1:500; Thermo Scientific, A21208), and donkey anti-sheep 647 (1:500; Thermo Scientific, A21448). Slides were mounted using Fluoromount-G^®^ (SouthernBiotech, 010001) and imaged on a Nikon A1R confocal microscope using NIS Elements (Nikon) in the Bioimaging and Analysis Facility at CCHMC. Immunofluorescent images were pseudocolored and the LUTs were adjusted using Fiji/ImageJ (NIH).

### Whole mount immunostaining

Organoids were washed with PBS and fixed in 4% PFA with 0.3% Triton X-100 at 4 °C for 60 min (small organoids < 300 µm) to 120 min (large organoids > 300 µm). The fixed samples were washed twice with PBS and once with distilled water. Tissues were then resuspended in 50% tetrahydrofuran (Thermo Scientific, 326970010) in water for 3 hours at room temperature, washed extensively with water, and permeabilized and blocked for 2 hours at room temperature with 5% donkey serum in PBST. Samples were incubated with primary antibodies for ∼36 hours at 4 °C, washed extensively and incubated with secondary antibodies for ∼36 hours at 4 °C. For imaging, organoids were resuspended in index matching solution made by diluting 13.13 g of urea (Sigma-Aldrich, U5378) and 25 g Histodenz™ (Sigma-Aldrich, D2158) in 11 mL of 0.02 M phosphate buffer. Samples were imaged on a Yokogawa W1 Spinning disk confocal attached to a Ti2 microscope (Nikon), with a 20x/0.95NA water immersion objective (Nikon) and an ORCA-Fusion BT camera (Hamamatsu). NIS Elements (Nikon) was used to make 3D rendering and movies of the organoid z-stack.

### Transmission electron microscopy imaging

Organoids were fixed in 3% glutaraldehyde overnight at 4C. Organoids were then washed using 0.1M sodium cacodylate buffer followed by a 1-h incubation using 4% osmiumtetroxide, washed and then dehydrated using 25–100% ethanol (series of dilutions), embedded using propylene oxide/Embed 812. Blocks were sectioned (100 nm) and stained with 2% uranyl acetate followed by lead citrate. Tissue was visualized using a transmission electron microscope (Hitachi HT-7800) with a digital camera (Biosprint 16).

### Luminal pancreatic enzymes quantification

Ten organoids were cut in two by using sharp disposable scalpels (Fisher Scientific, 16050117) under a stereo microscope. The luminal content of the ten organoids was collected and diluted in 350 µL of Krebs buffer. Krebs buffer (pH 7.4) contained 128 mM NaCl, 5mM KCl, 2.7 mM CaCl_2_·H_2_O, 1.2 mM MgSO_4_, 1 mM Na_2_HPO_4_, 1.2 mM KH_2_PO_4_, 5 mM NaHCO_3_, 10 mM HEPES, and 0.1% BSA. The luminal content was spun down at 10000 g for 4 min, and the supernatant was immediately frozen for later analysis. Trypsin activity was measured using a colorimetric assay (Abcam, ab102531), whereas CPA1 concentration was quantified using a human carboxypeptidase ELISA kit (Fisher Scientific, EH66RB), following the manufacturer’s protocols.

### Glucose stimulated insulin secretion

Ten organoids were cut in two by using sharp disposable scalpel (Fisher Scientific, 16050117) under a stereo microscope. The tissues were collected in a tube a gently pipetted up and down to remove debris before being placed in new Krebs buffer supplemented with 2.8 mM glucose to equilibrate to basal glucose for 2 hours under gentle shaking conditions (60 rpm). The insulin secretion test was performed by applying sequential Krebs solutions supplemented with 2.8 mM glucose for 30 min, 16.7 mM glucose for 30 min, and 30 mM KCl for 30 min. For the KCl, NaCl concentration was reduced to 104 mM to maintain adequate osmolarity. After each incubation, the supernatant was quickly centrifuged to remove potential cell debris and stored at -80°C. For analysis, the thawed samples were processed using an ELISA kit for human C-peptide (Alpco, 80-CPTHU-CH05, with standard curve) according to the manufacturer’s instructions. Absorbance (450 nm) was measured using a Synergy Neo2 Microplate reader (BioTek).

### In vivo transplantation of organoids

Organoids were all ectopically transplanted into the kidney capsule of NSG mice as previously described^35^. Briefly, one or two organoids were transplanted into the kidney subcapsular space. Engrafted organoids were harvested 12 weeks after transplantation, washed with PBS and fixed overnight in 4% PFA before being process for paraffin embedding and sectioning.

### Bulk RNA sequencing analysis

Cells and/or organoids were collected and digested in RA1 lysis buffer from the NucleoSpin RNA Kit (Macherey-Nagel, 740955.250) supplemented with beta-mercaptoethanol (Sigma) and stored in -80C. RNA extraction was performed using the above kit according to manufacturer protocol. Total RNA (>300 ng) was quality controlled, prepped and sequenced by the Integrated Genomics and Microbiome Sequencing Facility at CCHMC. Paired end 100 base pair polyA stranded RNA-sequencing was performed using NovaSeq X Plus (Illumina) to acquire 30 million reads per sample. To generate count matrix, raw FASTQ files were processed using the standard nf-co.re RNA-seq pipeline (https://nf-co.re/rnaseq/3.15.0) with the star salmon workflow to align to the human hg38 genome build. Analysis was performed in RStudio (v4.4.0). Differential gene expression analysis was performed using the DESeq2 R package (1.20.0) with a false discovery rate adjusted p-value <0.05 considered significant.

### Organoid Dissociation for single cell RNA-sequencing

Pancreatic organoids were washed with cold HBSS (ThermoFisher Scientific, 14175079) and finely minced using a scalpel. Minced organoids were enzymatically dissociated using enzymes from the Neural Tissue Dissociation Kit (Miltenyi Biotec, 130092628) prepared and added using manufacturer’s protocol and incubated at 37°C for ∼60 min with periodic fluxing through P1000 and P200 pipette tips. Cells where washed and resuspended in HBSS containing 1% BSA and Y-27632 (10 µM). All dissociated cells were filtered through BSA coated 40 μm cell strainers to remove undissociated tissues. A single cell suspension with >90% viability was confirmed using a cell counter (Bio-Rad).

### Single cell Library Preparation, RNA-sequencing, and Analysis

Single-cell RNA preparation and sequencing was performed by the Integrated Genomics and Microbiome Sequencing Facility at CCHMC. Briefly, for Batch 1, RNA from freshly dissociated single cells were isolated using Chromium X Next-GEM (10x Genomics) 3’v3.1 chemistry loading 12,800 cells to get an expected recovery rate of 8,000 with a multiplet rate of 6.2%. Batch 2, which consisted of D60 hPO+XLST only, utilized Chromium X GEX 3’v4 chemistry loading 21,750 cells to get an expected 15,000 cells with a multiplet rate of 6.0%. Samples from individual batches were sequenced at the same time using NovaSeq X Plus (Illumina) for approximately 30,000 reads per cell. Paired end FASTQ files were aligned to human hg38 genome using cellranger (Batch 1 v6.1.2 and Batch 2 v8.0.1) (10x Genomics) and samtools (Batch 1 v1.8.0 and Batch 2 v1.21.0). Filtered matrix, features and barcodes, were opened in R Studio (v4.4.0) and Seurat objects were made using standard Seurat (v5.3.0) pipeline requiring 3 and 200 minimum cells and features, respectively. Low quality cells were filtered based on nFeatures_RNA between 200-8000, nCounts > 1000, and mitochondrial percent < 20. Data was then normalized using standard NormalizeData and variable features were identified using vst selection method for nfeatures = 2000. The objects were scored based on known cell cycle genes to identify S, G2/M, G1 phases. Data was scaled regressing out mitochondrial and cell cycle signatures. Following scaling, data principal components were identified using RunPCA and then cells were grouped and visualized using FindNeighbors for 30 dimensions, FindClusters and RunUMAP functions in Seurat. Resolution of FindClusters was determined on an individual basis based on biologically relevant number of clusters. For both individual and integrated objects, count tables of metadata were used to determine cell proportions, FindAllMarkers with min.pct = 0.25 and logfc.threshold = 0.25 was used to determine cluster or sample enriched genes which was subsequently used for manual cluster annotations and genes of interest were visualized using FeaturePlot, DotPlot, and/or VlnPlot functions using ggplot2 (v3.5.2) package.

To infer potential cell-cell signaling between cell types, we utilized CellChat (v1.6.1) on individual objects. Cluster resolution was adjusted depending on detail required for signaling ranging from 0.01 (lpPO) to 0.025 (sPO) for low resolution (3-4 clusters) or 0.25 (lpPO) to 0.1 (sPO) for higher resolution (7-8 clusters). Clusters were renamed and made compatible using createCellChat function and underwent standard pipelines to identify ligands and receptors expression (>10 cells) in clusters to identify potential communication. Data was interpreted and visualized using a combination of tables for specific ligand and receptor details, as well as selected pathway heatmaps or circle plots focusing on difference between lpPO and sPO samples.

For integration of individual datasets into the whole hPO temporal atlas, individual objects were merged into a single object and went through a similar pipeline as described above. Object layers were integrated using Harmony (v1.2.3) integration. Object was further refined by filtering out nCount_RNA > 2000 and genes starting with MT-or RP. The final FindNeighbors used 1:30 dimensions, FindClusters with resolution 0.4, and RunUMAP with n.neighbors = 30 and min.dist = 0.4. To recluster specific cell populations, including all epithelium, exocrine, endocrine, or mesenchyme without endothelium, the whole atlas was subsetted based on relevant Seurat clusters and/or datasets before undergoing standard pipelines described above except for the RunHarmony function following PCA. Depending on reclustering, additional clusters potentially were removed, such as contaminating mesenchymal cells in endocrine recluster.

For human fetal tissues, we utilized published and unpublished 1^st^ and 2^nd^ trimester datasets. Annotated fetal pancreas or fetal endodermal tissue scRNA-seq Seurat objects were kindly provided by Dr. Julie Sneddon (UCSF) or Dr. Gray Camp (Max Planck Institute). Following UpdateSeuratObject from Seurat v3 to v5, objects went through pipeline described above, including cell cycle scoring, removal of MT-and RP genes, and filtering including nCounts_RNA > 2000 and Harmony integration. If not present, metadata including Organ, Tissue, Age, and Celltypes were added to fetal objects. To recluster specific cell populations, including all epithelium, exocrine, endocrine or mesenchyme, the whole atlas was subsetted based on relevant Seurat clusters as described above. Reclustered objects were further annotated with more detail. To integrate fetal pancreatic epithelium or mesenchyme with that of other fetal endodermal tissues, datasets were subsetted for common features present in both atlases RNA counts tables. Atlases were then merged and underwent harmony integration as described above. Following integration, the Liver sample was removed due to lack of sufficient number of cells. Furthermore, the epithelial atlas was also subclustered to generate endocrine-specific atlas, where the Esophagus sample was also removed due to lack of cells.

Using either human fetal pancreas atlas or combined fetal endodermal tissue atlas, we utilized LabelTransfer and/or MapQuery functions in Seurat to map fetal annotations, of either cell types or age metadata, onto integrated or individual hPO datasets, respectively. Layers were joined and datasets were subsetted for common features present in both atlases RNA counts tables. Reference anchors were identified using FindTransferAnchors function using 1:30 dimensions and PCA reduction and predictions were transferred onto query organoid dataset. In addition, we utilized the fetal datasets as a reference training dataset for a random forest machine learning algorithm to predict similarity utilizing the ranger (v0.17.0) package. Reference was downsampled based on reference metadata used to do the training. Downsampling for “Organ” level classification (whole atlas, all epithelium, all mesenchyme, endocrine) was 1000 cells for all except endocrine which was 35, “Celltypes” was 250 for exocrine, 50 for endocrine, and 200 for mesenchyme. For ranger random forest was trained using num.trees = 1000 trees and importance = “permutation”. Similarity predictions were visualized using UMAP or Heatmap of similarity matrix from 0 to 1 (most similar).

### Data Visualization

All figures were generated in InDesign (Adobe) and schematics were generated using Designer (Affinity). UMAPs, bulk RNA-seq heatmap, dot plots, volcano plots, feature plots, as well as cell-cell communication heatmaps were all generated in R Studio using Seurat and ggplot2 as described above. All other graphs and heatmaps were generated in Prism 10 (GraphPad).

### Data Accessibility

All scRNA-seq datasets generated in this study are available upon request from the authors. Published human fetal endodermal tissue scRNA-seq atlas used in this study are available from ArrayExpress under accession numbers: E-MTAB-10187, E-MTAB-10268, E-MTAB-8221, E-MTAB-9228, and E-MTAB-94. Human fetal pancreas scRNA-seq atlas used in this study was kindly provided by Dr. Julie Sneddon (UCSF). An alternative, single cell/nuclear RNA-seq human fetal dataset can be accessed from descartes.brotmanbaty.org. Previously published organoid scRNA-seq datasets can be accessed in the GEO under accession numbers: GSE214852, GSE240363, GSE254954. All scRNA-seq analysis in this study was performed in R using previously generated codes. This paper does not report original code. Additional information, original images or scripts needed for further analysis are available upon request.

## Acknowledgements

We thank the members of the Wells and Zorn laboratories for feedback. We thank Dr. Elliott Brooks for coordinating the scRNA-sequencing data transfer. We acknowledge core support from the CCHMC, including the Pluripotent Stem Cell Facility (RRID:SCR_022634), Single Cell Genomics Facility (RRID:SCR_022653), Genomics Sequencing Facility (RRID:SCR_022630), Integrated Pathology Research Facility (RRID:SCR_022637), Research Flow Cytometry Facility (RRID:SCR_022635), Bio-Imaging and Analysis Facility (RRID:SCR_022628) and Veterinary Services Facility. This research was supported by grants from the Allen Foundation and National Institutes of Health R01-CA272903 and P01-HD093363 to J.M.W. This work was also supported by grants to J.B.S. from the NIH/NIDDK (grants R01DK118421 and R01DK144425). J.A.B. was supported by Schmidt Science Fellows, in partnership with the Rhodes Trust. Support was also received from the Digestive Disease Research Center (P30-DK078392).

## Contributions

J.A.B. and J.M.W. conceived the study and experimental design, interpreted data and wrote the manuscript. D.O.K. and J.A.B. performed bioinformatic analysis. P.T. and J.A.B. performed the experiments. L.D. and M.K. provided fetal sections and staining. J.B.S. provided fetal single cell RNA sequencing data. All authors contributed to the editing of the manuscript.

---

**Extended Data Figure 1:**
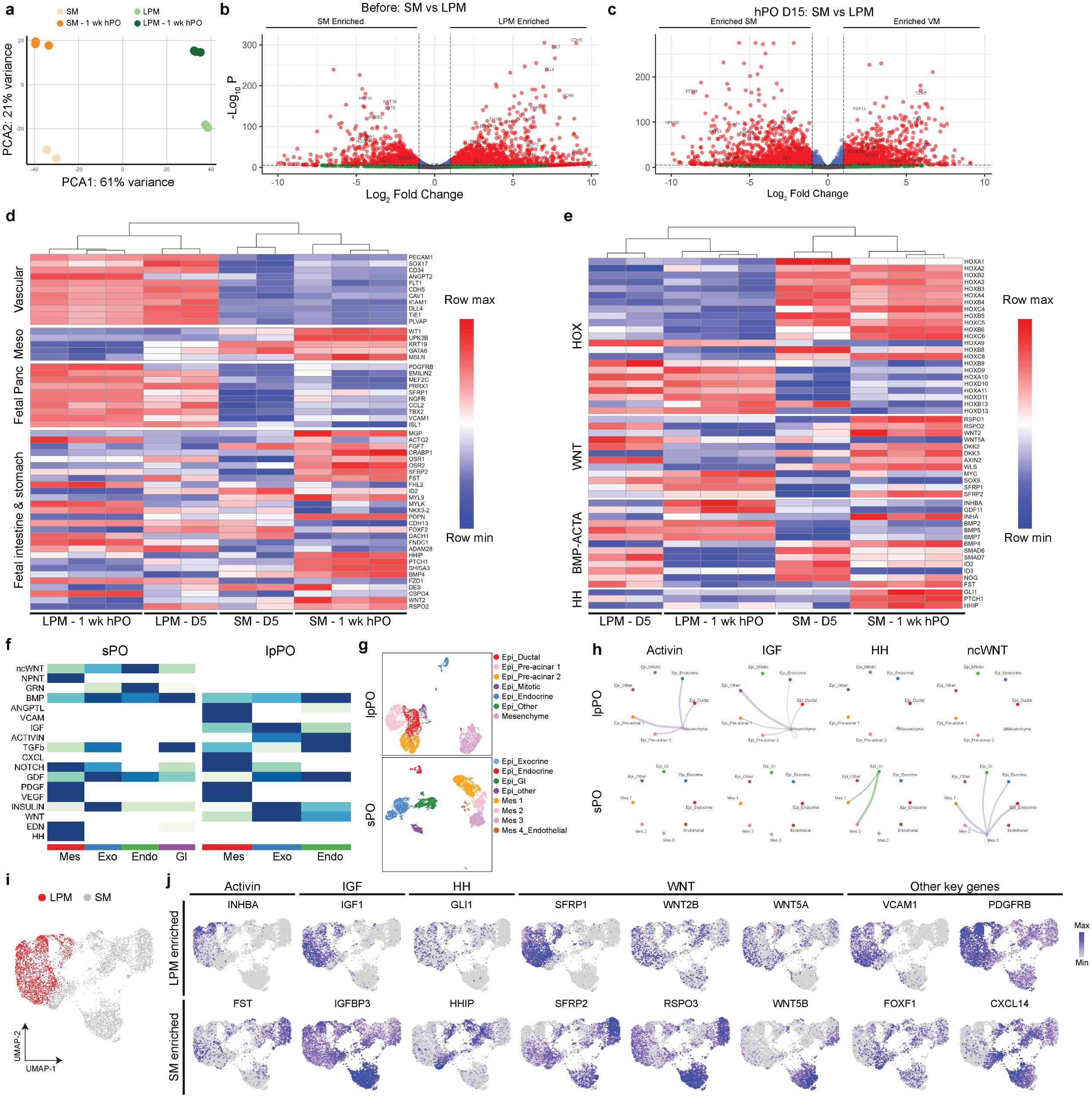
Comparison of mesenchyme identity. **a**, PCA of splanchnic mesenchyme and lateral plate mesenchyme before recombination or sorted after 1 week of culture within the hPOs. **b-c**, Volcano plots showing key differentially expressed genes between the splanchnic and lateral plate mesenchyme before (**b**) or after (**c**) recombination. **d-e**, Heatmap of differentially expressed genes between splanchnic and lateral plate mesenchyme before and after recombination. **f**, Predicted incoming cell-cell crosstalk between major cell types in sPO and lpPO as determined by CellChat. **g**, UMAP projections of sPO and lpPO, color-coded based on cell type annotations. **h**, Cell-cell communication inference for selected statistically significant pathways between sPO and lpPO. **i**, UMAP of scRNA-seq datasets for lpPO and sPO mesenchyme at day 25, colored by experimental conditions. **j**, Feature plots showing expression levels of indicated genes.

**Extended Data Figure 2:**
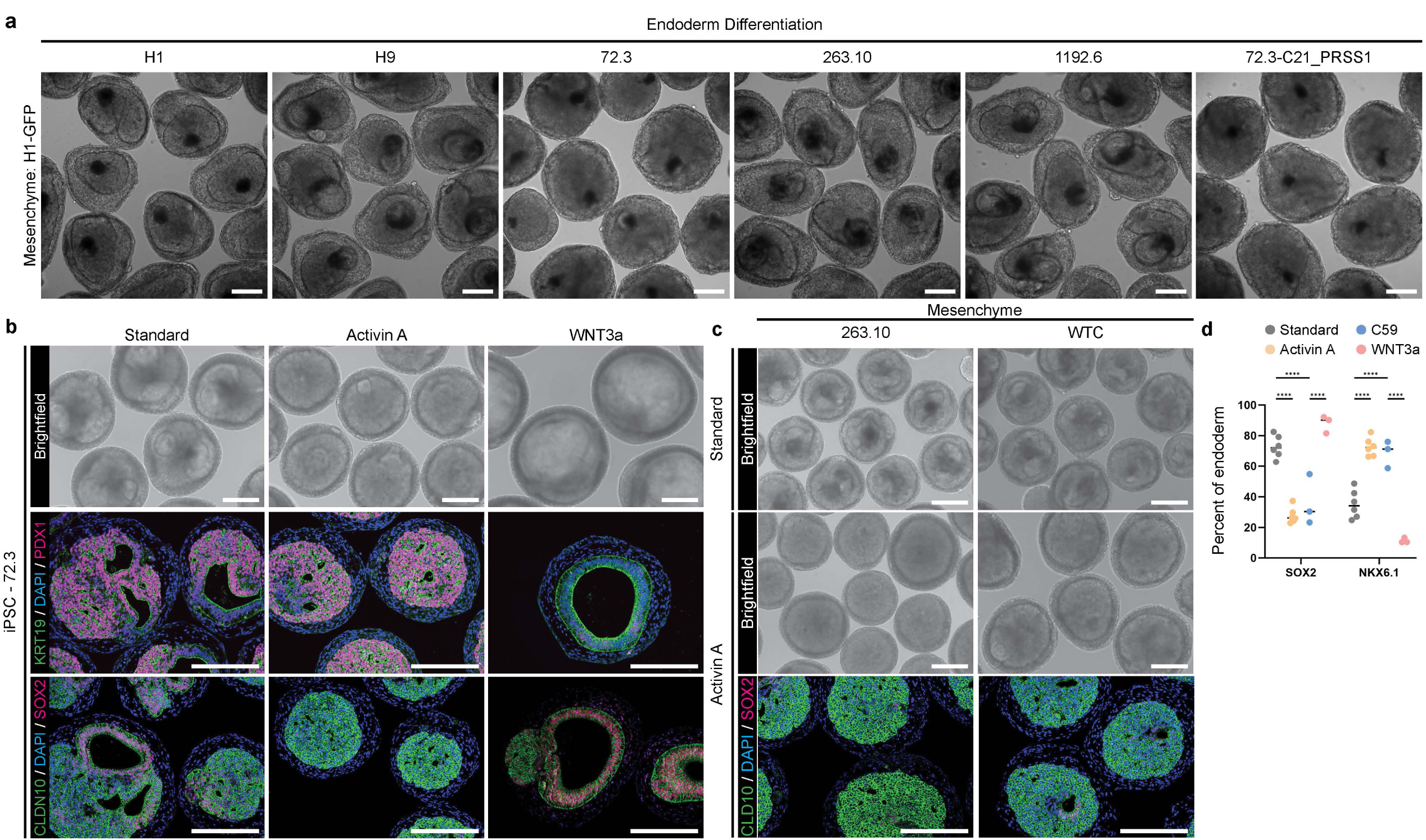
Protocol robustness. **a**, Representative brightfield images of day 25 organoids recombined with H1-gfp LPM and PFG from different cell lines. Scale bars, 200 µm. **b-c**, Representative brightfield (top) and immunostaining (bottom) images of day 25 hPO made from the iPSC line 72.3 (**b**) or 263.10 and WTC (**c**). Scale bars, 200 µm. **d**, Flow cytometry analysis showing the percent of PFG derived cells that express SOX2 and NKX6.1 in function of the culture conditions. The data presented is for the iPSC line 72.3. Mean and standard deviation are shown for 6 (Standard, Activin A) or 3 (WNT3a, C59) independent differentiations. ***p < 0.001 and ****p < 0.0001, determined by two-way ANOVA with Sidak’s multiple comparisons test.

**Extended Data Figure 3:**
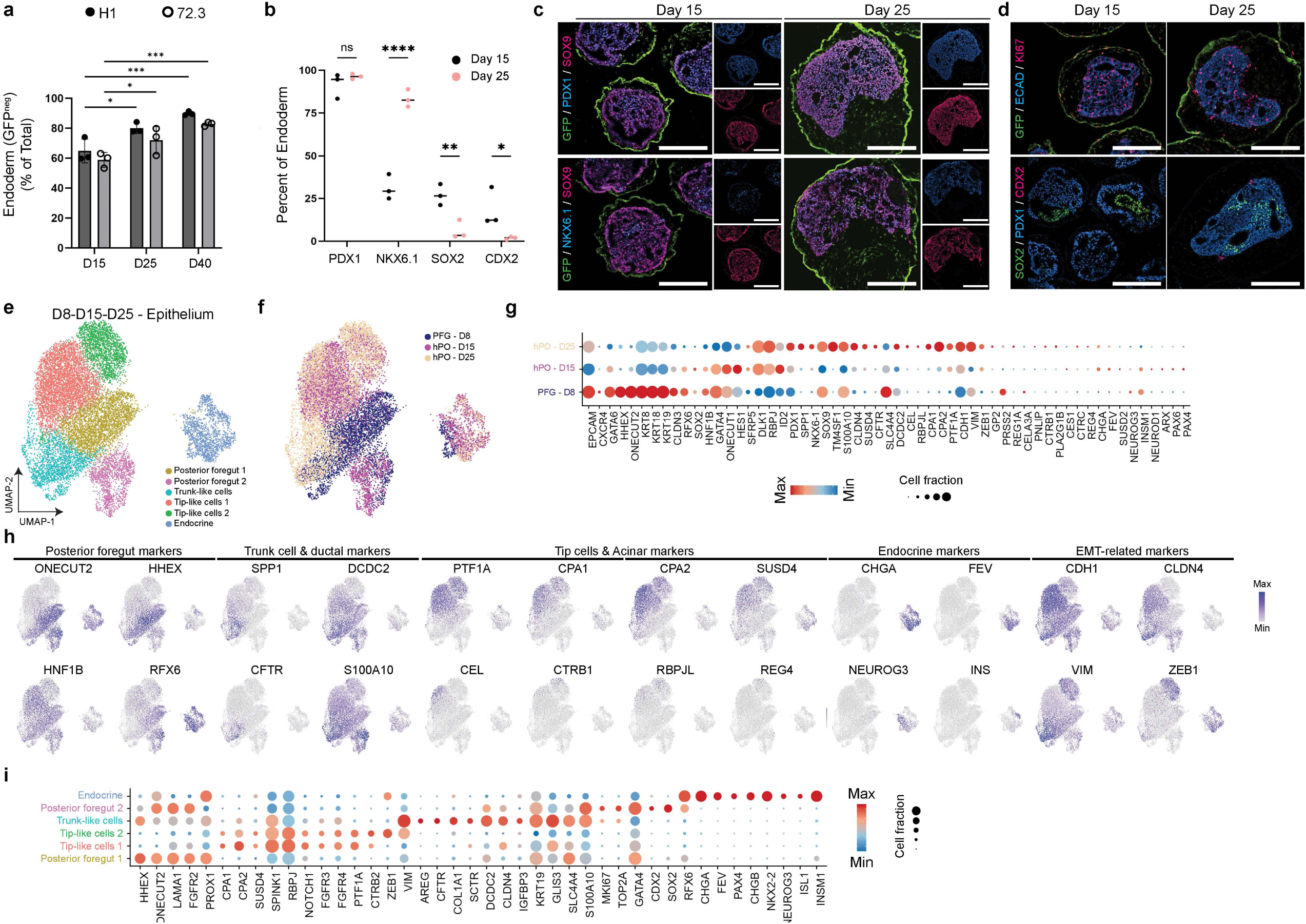
Characterization of pancreatic progenitors. **a**, Flow cytometry analysis showing the percentage of PFG derived cells (GFP^neg^) expressing different transcription factors for day 15 and 25 hPOs. Mean and standard deviation are shown for 3 independent differentiations. *p < 0.05 and ***p < 0.001, determined by two-way ANOVA with Tukey’s multiple comparisons test. **b**, Flow cytometry analysis showing the percentage of PFG derived cells (GFP^neg^) in function of organoid development. Mean and standard deviation are shown for 3 independent differentiations. *p < 0.05, **p < 0.01 and ****p < 0.0001, determined by two-way ANOVA with Sidak’s multiple comparisons test. **c**, Representative immunostaining images of day 15 and 25 hPOs showing merged pseudocolor (left) and single color (right). Scale bars, 200 µm. **d**, Representative immunostaining images of day 15 and 25 hPOs showing proliferation (Ki67) and endoderm patterning (SOX2, CDX2, PDX1). Scale bars, 200 µm. **e-f**, UMAP projections of epithelium from PFG and early organoids (day 15 and day 25), with cell identity annotated by clusters (**e**) or timepoint (**f**). **g**, Dot plot showing the expression of key marker genes across organoid timepoint. Dot sizes and colors indicate proportions of cells expressing the corresponding genes and their averaged scaled values of log-transformed expression, respectively. **h**, Feature plots showing expression levels of indicated genes. **i**, Dot plot showing the expression of key marker genes across clusters. Dot sizes and colors indicate proportions of cells expressing the corresponding genes and their averaged scaled values of log-transformed expression, respectively.

**Extended Data Figure 4: s.**
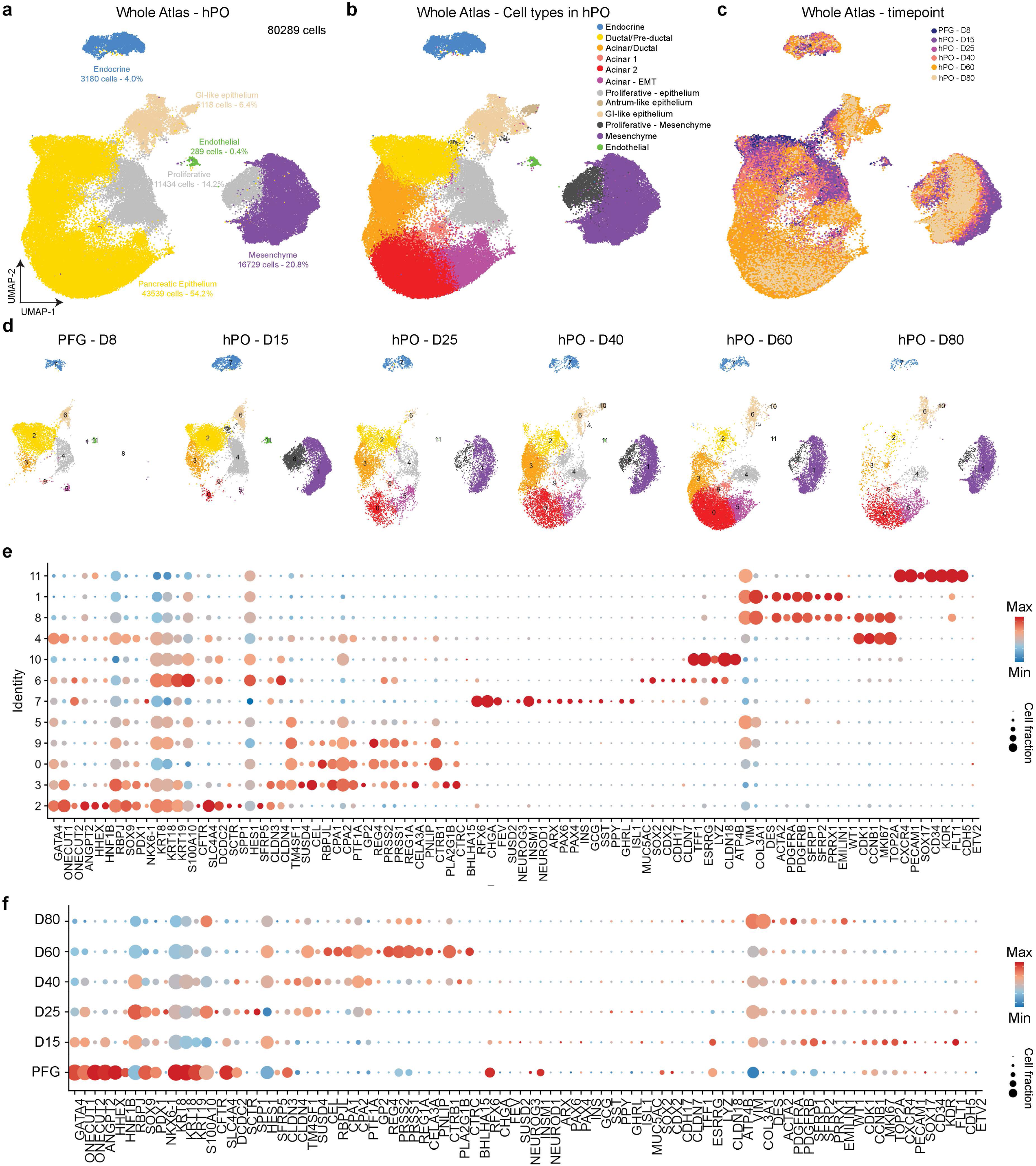
c-RNAseq hPO atlas. **a-c**, UMAP projections of whole hPO atlas, color-coded based on broad cell type categories (**a**), specific cell types (**b**) and timepoint (**c**). **d**, UMAP projections of hPO atlas, split between the different organoid development timecourse. **e-f**, Dot plot showing the expression of key marker genes across clusters (**e**) and organoid timepoint (**f**). Dot sizes and colors indicate proportions of cells expressing the corresponding genes and their averaged scaled values of log-transformed expression, respectively.

**Extended Data Figure 5:**
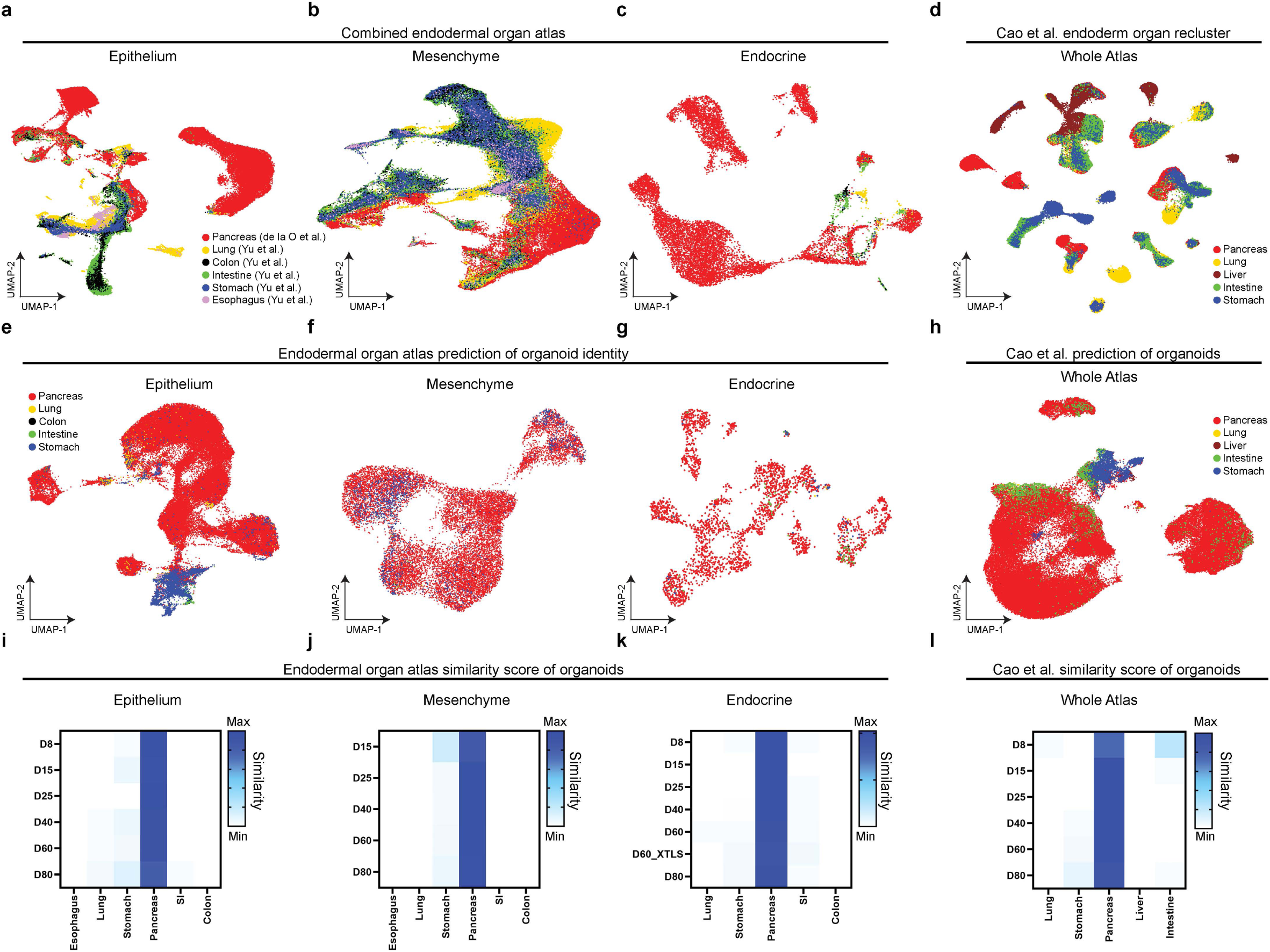
Organ specificity of hPO. **a-c**, UMAP of combined endodermal organ atlas of epithelial (**a**), mesenchymal (**b**) or endocrine (**c**) cells, color-coded based on the different organs^25,34^. **d**, UMAP of the *Cao et al*. atlas^24^, reclustered to include only endodermal organs. **e-h**, UMAP of scRNA-seq data showing identity prediction of the organoids against the epithelial, mesenchymal, endocrine or whole endodermal atlas. **i-l**, Similarity score between specific cell type within the organoids (epithelial, mesenchymal and endocrine) and different atlas, obtained using a machine learning algorithm for identity prediction.

**Extended Data Figure 6:**
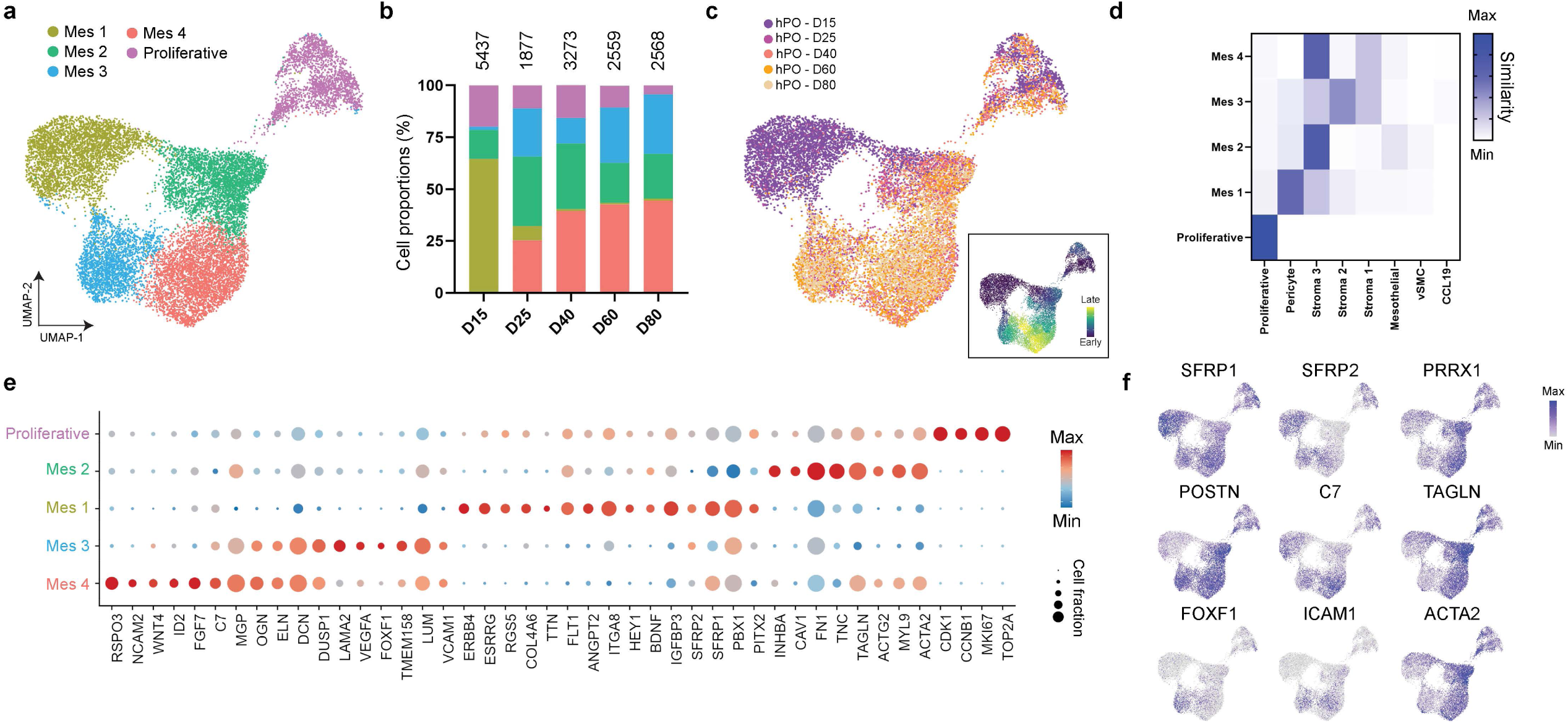
Characterization of hPO mesenchyme. **a**, UMAP projections of mesenchymal cells subsetted from the hPO atlas, with cell identity annotated by clusters. **b**, Bar graph showing the cell proportions for each cell type throughout the differentiation, color-coded using the cell clusters defined in **a**. **c**, UMAP projections of mesenchymal cells from the hPO atlas, color-coded by timepoints. **d**, Similarity score between mesenchymal clusters within the organoids (y axis) and specific cell types found in the fetal pancreatic mesenchyme atlas (x axis)^25^, obtained using a machine learning algorithm for identity prediction. **e**, Dot plot showing the expression of key marker genes across clusters. Dot sizes and colors indicate proportions of cells expressing the corresponding genes and their averaged scaled values of log-transformed expression, respectively. **f**, Feature plots showing expression levels of indicated genes.

**Extended Data Figure 7:**
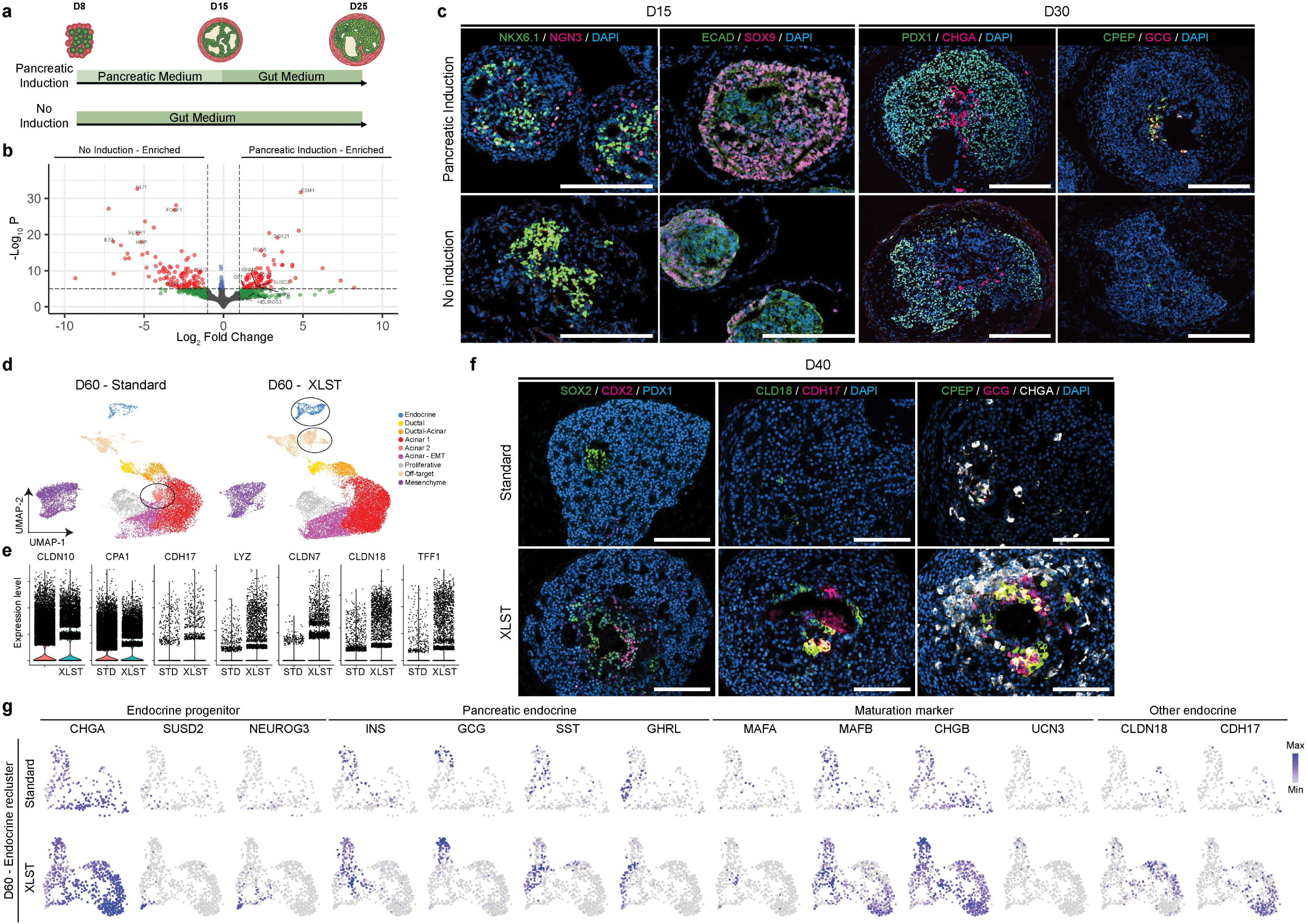
Effect of endocrine induction on hPO differentiation. **a**, Schematic summary of the differentiation protocol with or without pancreatic induction during the first week of culture. **b**, Volcano plots showing key differentially expressed genes between the organoids cultured with pancreatic induction medium or without. **c**, Representative immunostaining images of organoids at day 15 or 30 in function of the culture conditions. Scale bars, 100 µm. **d**, UMAP projections of hPOs treated with or without endocrine induction cocktail (XLTS), color-coded based on cell type annotations. **e**, Violin plot of key genes showing similar or different expression patterns between the two organoid protocols. **f**, Representative immunostaining of day 40 hPOs showing key differences in protein expression of genes related to patterning (left, middle) or endocrine differentiation (right). Scale bars, 100 µm. **g**, Feature plots showing expression levels of indicated genes.

